# Developmental trajectories and differences in functional brain network properties of preterm and at-term neonates

**DOI:** 10.1101/2023.11.24.568614

**Authors:** N López-Guerrero, S Alcauter

**Affiliations:** Instituto de Neurobiología, Universidad Nacional Autónoma de México, Querétaro, México

**Keywords:** Brain network, Neonates, Development, Preterm, Graph theory

## Abstract

Premature infants, born before 37 weeks of gestation can have alterations in neurodevelopment and cognition, even when no anatomical lesions are evident. Resting-state functional neuroimaging of naturally sleeping babies has shown altered connectivity patterns, but there is limited evidence on the developmental trajectories of functional organization in preterm neonates. By using a large dataset (n=597) from the developing Human Connectome Project, we explored the differences in graph theory properties between at-term and preterm neonates at term-equivalent age, considering the age subgroups proposed by the World Health Organization for premature birth. Leveraging the longitudinal follow-up for some preterm participants, we characterized the developmental trajectories for preterm and at-term neonates. We found significant differences between groups in connectivity strength, clustering coefficient, characteristic path length and global efficiency. Specifically, at term-equivalent ages, higher connectivity, clustering coefficient and efficiency are identified for neonates born at later postmenstrual ages. Similarly, the characteristic path length showed the inverse pattern. These results were consistent for a variety of connectivity thresholds at both the global (whole brain) and local level (brain regions). The brain regions with the greatest differences between groups include primary sensory and motor regions and the precuneus which may relate to the risk factors for sensorimotor and behavioral deficits associated with premature birth. Our results also show non-linear developmental trajectories for premature neonates, but decreased integration and segregation even at term-equivalent age. Overall, our results confirm altered functional connectivity, integration and segregation properties of the premature brain despite showing rapid maturation after birth.

## Introduction

According to the World Health Organization (WHO), preterm birth is a live birth that occurs before 37 weeks of gestational age (GA). The WHO classifies preterm birth according to weeks of gestation (WG): extremely preterm, born < 28 WG; very preterm, between 28 to 32 WG; and moderate to late preterm: 32 to 37 WG (*Preterm Birth*, 2022). In 2020, an estimated 13.4 million preterm infants were born worldwide, a slightly lower rate compared to 2010, where it was 13.8 million. However, over the last decade, the global annual rate of reduction is estimated at -0.14% and very few countries have achieved this reduction (Geneva: World Health & Organization, 2023; Ohuma et al., 2023). Mortality and morbidity risks increase as a function of the degree of prematurity, including the risk for neurodevelopmental deficits (Perin et al., 2022; World Health Organization, 2012,2023).

The developmental consequences of premature birth can be neurological, cognitive, behavioral, and emotional, even when no anatomical brain lesions are evident; the severity increases with decreasing gestational age at birth, and the sequelae can last into adulthood (Larsen et al., 2022; Ream & Lehwald, 2018; Rogers et al., 2018). A premature infant is likely to survive from 25 weeks of GA at a critical time of development and vulnerability to the extrauterine environment, which could influence the child’s later development. Among the most frequent alterations are attention deficit hyperactivity disorder (ADHD), depression, anxiety, autism spectrum disorder (ASD), cerebral palsy (CP), antisocial personality, auditory and visual deficits, and an increased risk for cognitive, sensory, motor and language deficits (Hee Chung et al., 2020)

In recent years, neuroimaging methods have allowed the exploration of the brain *in vivo*; in particular, resting state functional Magnetic Resonance Imaging (rsfMRI) has made it possible to study the functional organization of the brain in fetus, preterm and term neonates (Smyser et al., 2016; Smyser & Neil, 2015; Thomason et al., 2015). Most rsfMRI studies in preterm infants have focused on characterizing the main brain networks in the resting state, showing that the primary networks are similar to those of adults. In contrast, there appears to be no complete precursor of the default mode network (DMN) in premature infants at term-equivalent age (Doria et al., 2010; Eyre et al., 2021). When characterizing the brain as a complex network, empirical evidence shows that from 20 WG, the fetal brain follows a modular organization with the centers being densely interconnected, forming a rich club organization; with modules representing the areas that will give rise to the motor, visual, auditory, and the DMN, among others (Thomason et al., 2014). By 31 WG, the network organization starts to become more efficient, specialized, and integrated with a stronger connection between modules than towards the interior of modules; there is a decrease in local efficiency as age increases the clustering coefficient and shortest path length (De Asis-Cruz et al., 2021; Thomason et al., 2014; Turk et al., 2019). Modular structure, clustering coefficient, and shortest path length continuously change during the neonatal period (Gao et al., 2011). These network properties are also observed in premature infants at term-equivalent age, showing however, reduced rich club organization, less segregated networks, less integration and reduced global efficiency (Bouyssi-Kobar et al., 2019; Scheinost et al., 2016). Nevertheless, there is limited documentation regarding the functional organization in preterm neonates, and the characterization of developmental trajectories for this population has been challenging mainly due to small sample sizes.

In this study, leveraging a large public dataset, we examined whether premature infants, at term-equivalent age, exhibit brain functional organization similar to that of at-term neonates, both at the global and local levels. By using data from the Developing Human Connectome Project (Hughes et al., 2017), including longitudinal follow ups for some premature infants, we delved into the developmental trajectories of these network properties in both premature and at-term neonates.

## Methods

### Participants

We analyzed 418 datasets from the developing Human Connectome Project (Hughes et al., 2017, third open access data; see Supplementary Material for the list of IDs used here), corresponding to complete preprocessed structural and functional brain imaging, acquired between 23-44 weeks postmenstrual age (PMA), with a radiologist score between 1 and 2, *i.e.* with no significant incidental findings (see below). We extracted two samples; the first cross-sectional sample consisted of 393 scans taken at term-equivalent age, 336 considered at-term with a mean GA of 40.07 weeks (IQR=1.86), scanned at an average PMA of 41.55 weeks (IQR=2.57) and 57 preterms with a mean GA of 32.04 weeks (IQR=5.43) scanned at an average PMA of 41.21 weeks (IQR=2.15). The second, consisted of a longitudinal sample (50 fMRI/25 infants), two scans per subject, the first at a few days after birth (average PMA = 33.96 weeks, IQR=2.14) and the second at the term-equivalent age (average PMA = 41.49 weeks (IQR=1.72)).

### Image acquisition and preprocessing

The families of the infants were recruited at St Thomas’ Hospital, London, and the images were taken at the Evelina Newborn Imaging Centre, Centre for the Developing Brain, King’s College London, United Kingdom. The study was approved by the UK Health Research Authority Ethics Committee (14/LO/1169) and written informed consent was obtained from the parents to collect and disclose the images, as described in Hughes et al. (2017). Images were obtained in natural sleep in the MRI room located within the neonatal intensive care unit using a Philips Achieva 3T system (using a modified R3.2.2 software), with a 32-channel receiver head coil, acoustic hood, optimized transport system, positioning devices and hearing protection specially designed for this population, allowing optimum head position control and increased comfort. Infants were fed, swaddled, and placed in a vacuum sheath; scans were supervised by a neonatal nurse or pediatrician who monitored oxygen saturation, heart rate, and body temperature throughout the study (Hughes et al., 2017).

The fMRIs were acquired using a 9x accelerated multiband (MB) echo-planar imaging sequence over 15 minutes and 3 seconds, with repetition time (TR)= 392 ms, echo time (TE) = 38 ms, voxel size = 2.15 x 2.15 x 2.15 mm^3^, flip angle = 34°, 45 slices and a total of 2300 volumes. T1-weighted images had a reconstructed voxel size = 0.8 x 0.8 x 0.8 mm^3^, a field of view = 145 x 122 x 100 mm^3^, TR = 4795 ms and T2-weighted images had reconstructed voxel size = 0.8 x 0.8 x 0.8 mm^3^, a field of view = 145 x 145 x 108 mm, TR = 12 s, TE = 156 ms (Edwards et al., 2022). A radiologist evaluated all scans with scores from 1 to 5, with 4 and 5 showing significant lesions in the white matter, cortex, cerebellum, or basal ganglia. The most common incidentals included punctate white matter lesions (PWML) (12%) and caudothalamic subependymal cysts (10%) (Carney et al., 2021).

### Data processing

Individual fMRI were preprocessed with a pipeline made explicitly for the dHCP neonatal data to minimize artifacts in the BOLD signal (for details, refer to Fitzgibbon et al., 2020). Summarizing, preprocessing included field distortion, intra- and inter-volume motion corrections, removal of structured noise and cardiorespiratory artifacts, and regression of physiological noise and motion related estimates, including frame-wise displacement and the spatial standard deviation of successive difference images DVARS (Power et al., 2014). Finally, fMRI datasets were registered to the Infant Brain Atlas from Neonates brain template using linear and nonlinear transformations. We further delimited the fMRI datasets exclusively to gray matter voxels according to the dHCP atlas.

### Network measures

A graph is a non-empty set of nodes or vertices and a set of edges or links between pairs of nodes. The brain network may be represented as a graph where the nodes are defined brain regions and the links, in this case, are the value of functional connectivity between them, defined as the correlation between their BOLD signals (Friston, 1994). For each subject, we estimated the correlation between the BOLD time-series of all possible pairs of the 90 regions within the neonate AAL atlas (Shi et al., 2011); the resulting matrix represents the brain functional network for each subject. To minimize the number of variables and keep the number of connections constant between subjects and groups, the correlation matrices were thresholded and a fixed proportion of the strongest connections was kept. To avoid biases by selecting a single threshold, various proportion thresholds (also known as cost values) were tested from 0.05 to 0.50 in steps of 0.05 (that is, keeping 5 to 50 % of the strongest connections in steps of 5 %). From these thresholded matrices, we estimated network properties as described below.

Network properties can be studied at the local (nodes) or global (the whole network) level. To compare the properties of the brain networks of preterm and at-term infants, we used four properties: 1) node strength (*S*), 2) clustering coefficient (*CC*), 3) length of the shortest path (*d*) and 4) global efficiency (*E*_*global*_) using the Brain Connectivity Toolbox (Rubinov & Sporns, 2010). Detailed definitions are provided below, according to Rubinov & Sporns (2010).

Through these parameters, we can study two fundamental properties to understand brain functional organization: segregation, which refers to specialized densely interconnected groups of brain regions, and integration, which estimates the potential to share information globally. Higher *CC* indicates a segregated network and shorter *l* and higher *GE* indicates high integration.

The node strength (*S*), also known as weighted degree or connectivity strength, is defined as the sum of the weights *w* of all edges connected to a node *i*:

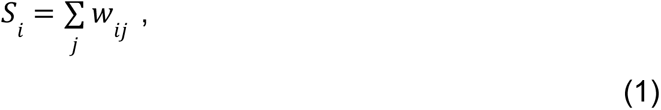

The clustering coefficient (*CC*) measures the average probability that two neighbors of node *i* are also connected, i.e. the proportion of triangles around a single node at the network level, is defined as the average value of *C_i_* over all nodes (*n*) in the network where *N* is the total number of nodes (Watts y Strogatz, 1998):

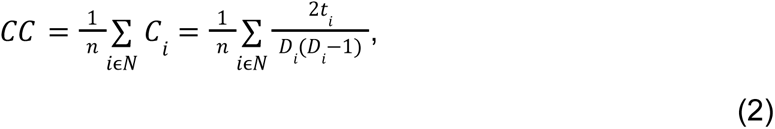

where the degree of connectivity of a node *i* is denoted as *Di* and *t*_*i*_ is the number of triangles in the graph that includes all the nodes that node *i* forms with all the other nodes in *N*.

The average length of the shortest path (*d*) is also called the characteristic path length, *d*_*ij*_ is defined as the length of the shortest path going from nodes *i* to *j* (Watts & Strogatz, 1998):

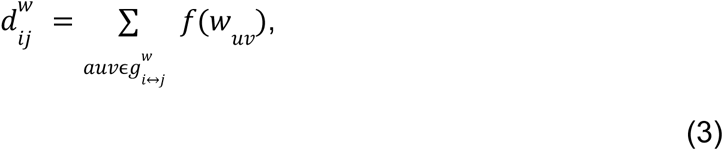

where *f* is a mapping from weight to length and 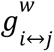 is the shortest weighted path between nodes *i* and *j*.

The global efficiency (*E*_*global*_) is the average of the inverse of the shortest path length from a node to all other nodes *i* other than *j*, measures the efficiency of parallel communication when all nodes exchange information with each other at the same time (Liu):

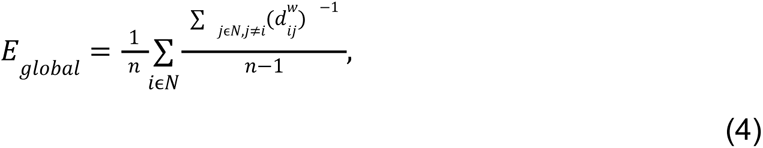

where 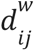 is the shortest path length between node *i* and node *j*.

### Division of groups

Figure 1 shows the total sample of neonates that we grouped based on the classification proposed by the WHO. Specifically, preterm infants are classified according to PMA as follows:

- Extremely preterm: < 28 WG
- Very preterm: 28 to 32 WG
- Moderate to late preterm: 32 to 37 WG

**Figure 1.**
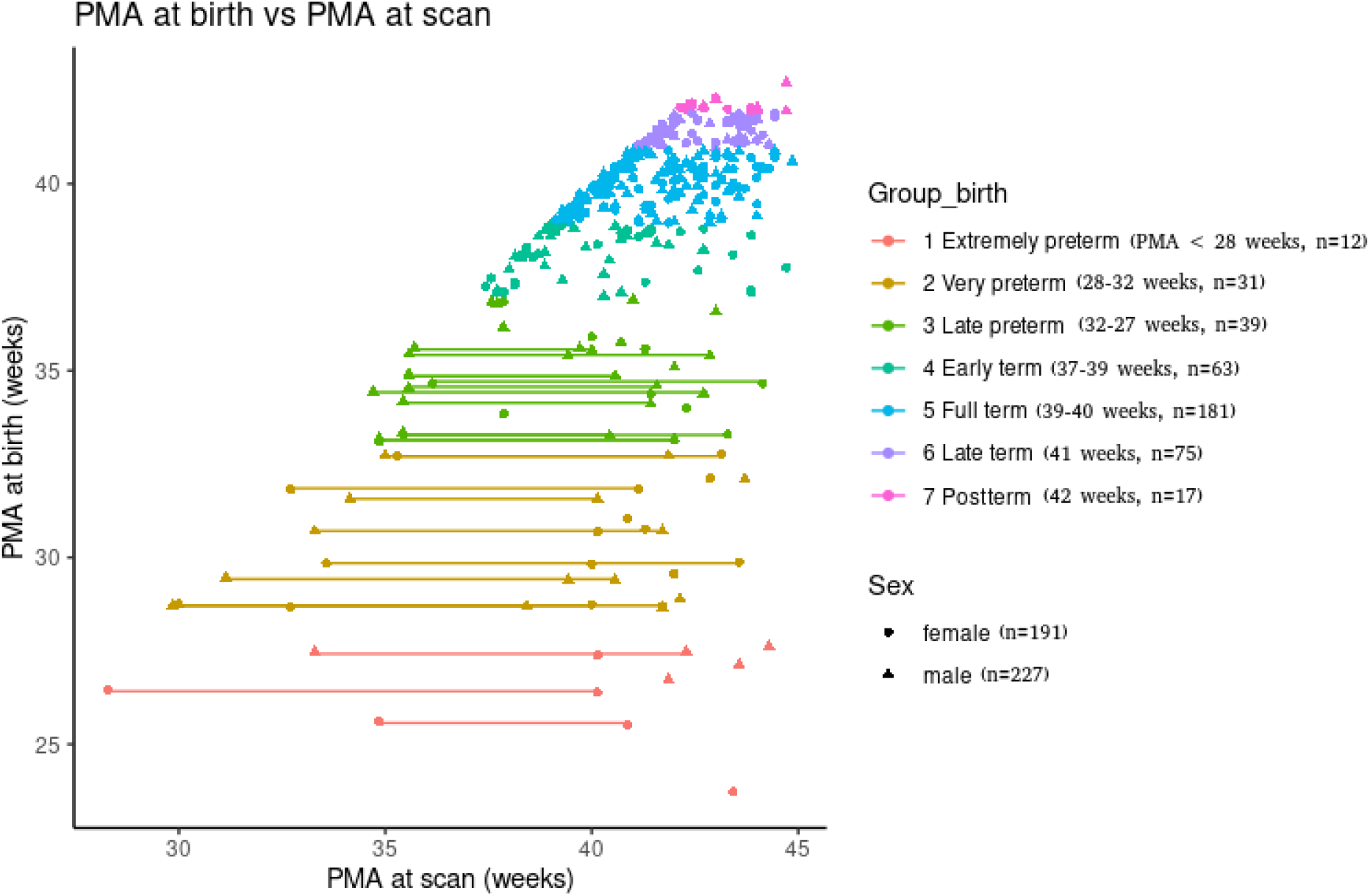
PMA of the sample. PMA at birth and at scan for the sample here explored (lines represent longitudinal data for the same participant).

In at-term infants, variability has been demonstrated in a variety of domains depending on the gestational week of delivery within a 5-week range - e.g.respiratory morbidity - and therefore, the American College of Obstetricians and Gynecologists and the Society for Maternal-Fetal Medicine (2012) proposed a more detailed classification that we also adopted for grouping the at-term neonates. Specifically, the following grouping was implemented:

- Early term: 37 to 38 WG
- Full term: 39 to 40 WG
- Late term: 41 WG
- Postterm: > 42 WG

To characterize the developmental trajectories of the graph theory properties here explored, linear, quadratic, and log-linear mixed models were constructed with gestational age at scan as an independent fixed-effect variable. Random effects were added for the intercept and subject ID. Significance was defined as p < 0.05, and the model with the lowest Akaike Information Criterion (AIC) was selected as the best model to describe the data.

## Results

First, we observed the connectivity values for each group as histograms. Figure 2 shows the distribution of the correlation values by group, with the average values between 0 and 0.5 for all groups. However, the average values are lower for the premature groups, increasing with PMA at birth.

**Figure 2.**
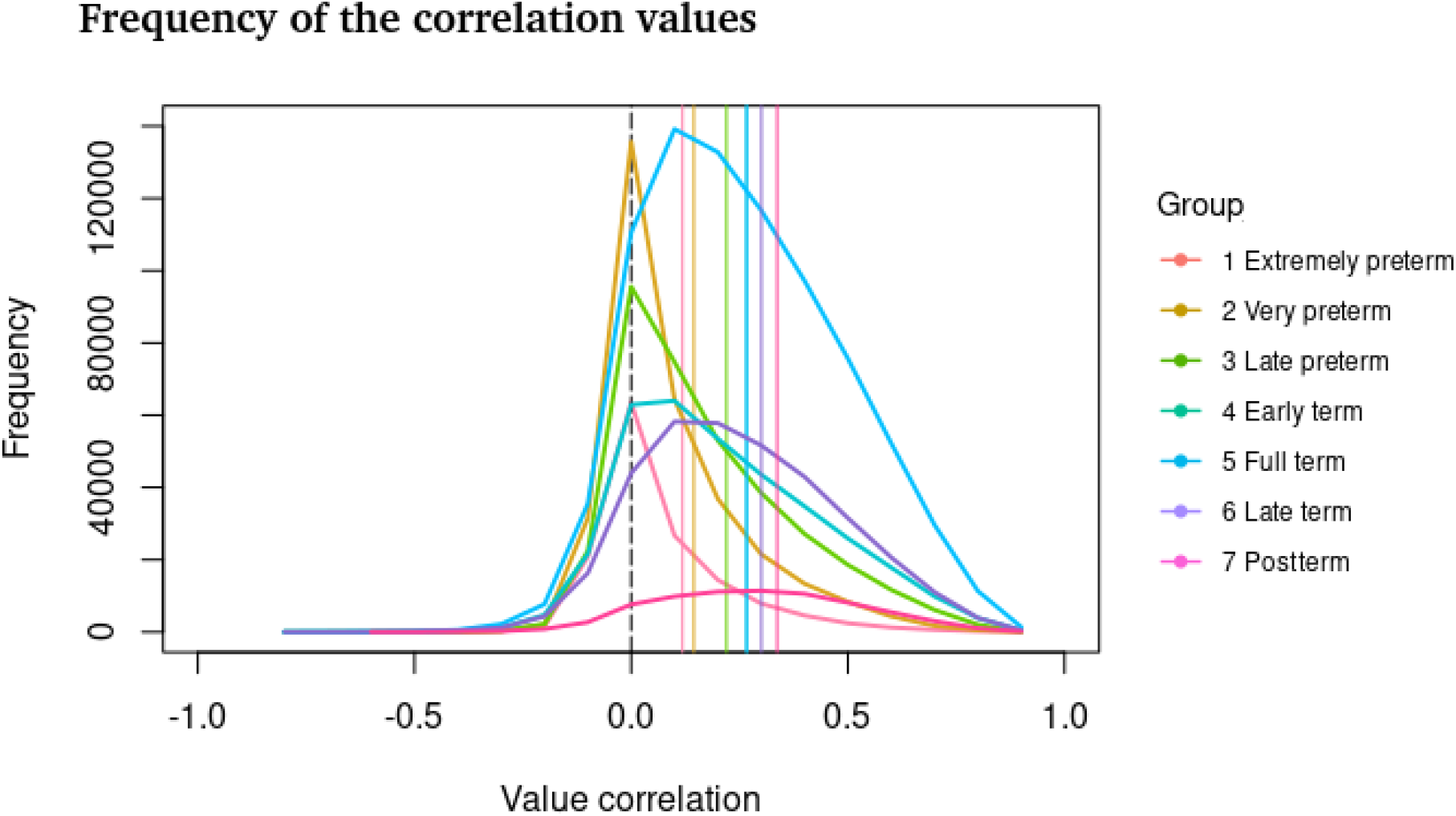
Frequency of the correlation values per group. The average correlation values increase with the PMA at birth (vertical lines indicate the average for each group).

When contrasting the network properties between groups, we observed that all cost values show significant differences between groups for all graph theory measures (ANCOVA tests; P<0.05, FDR corrected, Figure 3).

**Figure 3.**
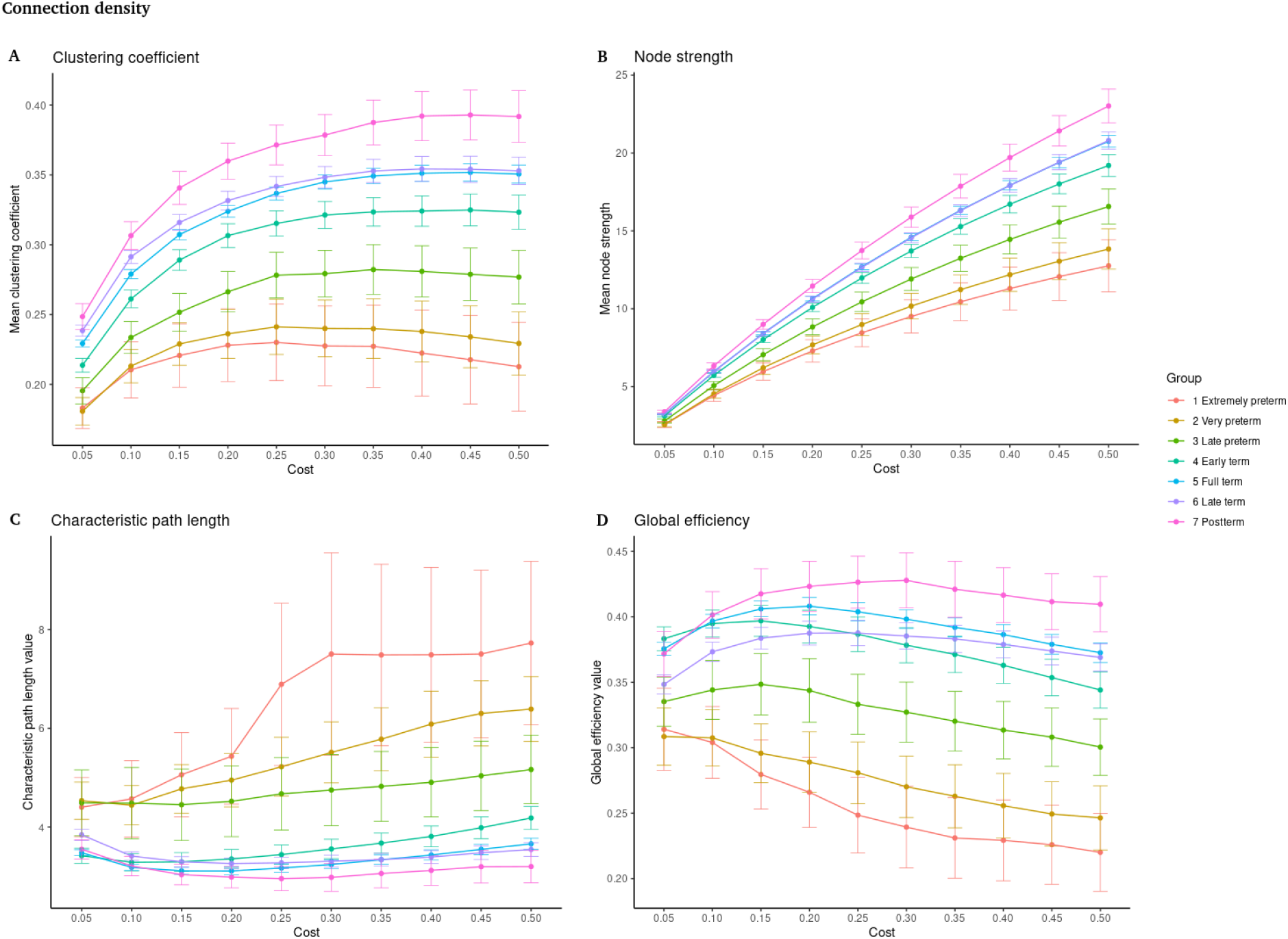
Graph theory properties as a function of the connection density. A ) Clustering coefficient. B) Node strength. C) Characteristic path length. D) Global efficiency. Significant differences are shown for all costs (percent of total connections) between all groups (P<0.05, FDR corrected). On the x-axis, the cost values are shown from 0.05 to 0.50 in steps of 0.05. On the y-axis, the values of the graph theory measures are shown.

For simplicity, we further detail differences between groups considering only the cost value of 0.10 (strongest 10% of all the connections).

### 1. Cross-sectional dataset, differences at term-equivalent age

To investigate whether preterm infants scanned at term-equivalent age exhibited the same global network properties as infants born at-term, a sample of 393 infants acquired between 37-44 weeks of postmenstrual age (PMA) and without radiological signs of white matter lesions was explored (Figure 4).

**Figure 4.**
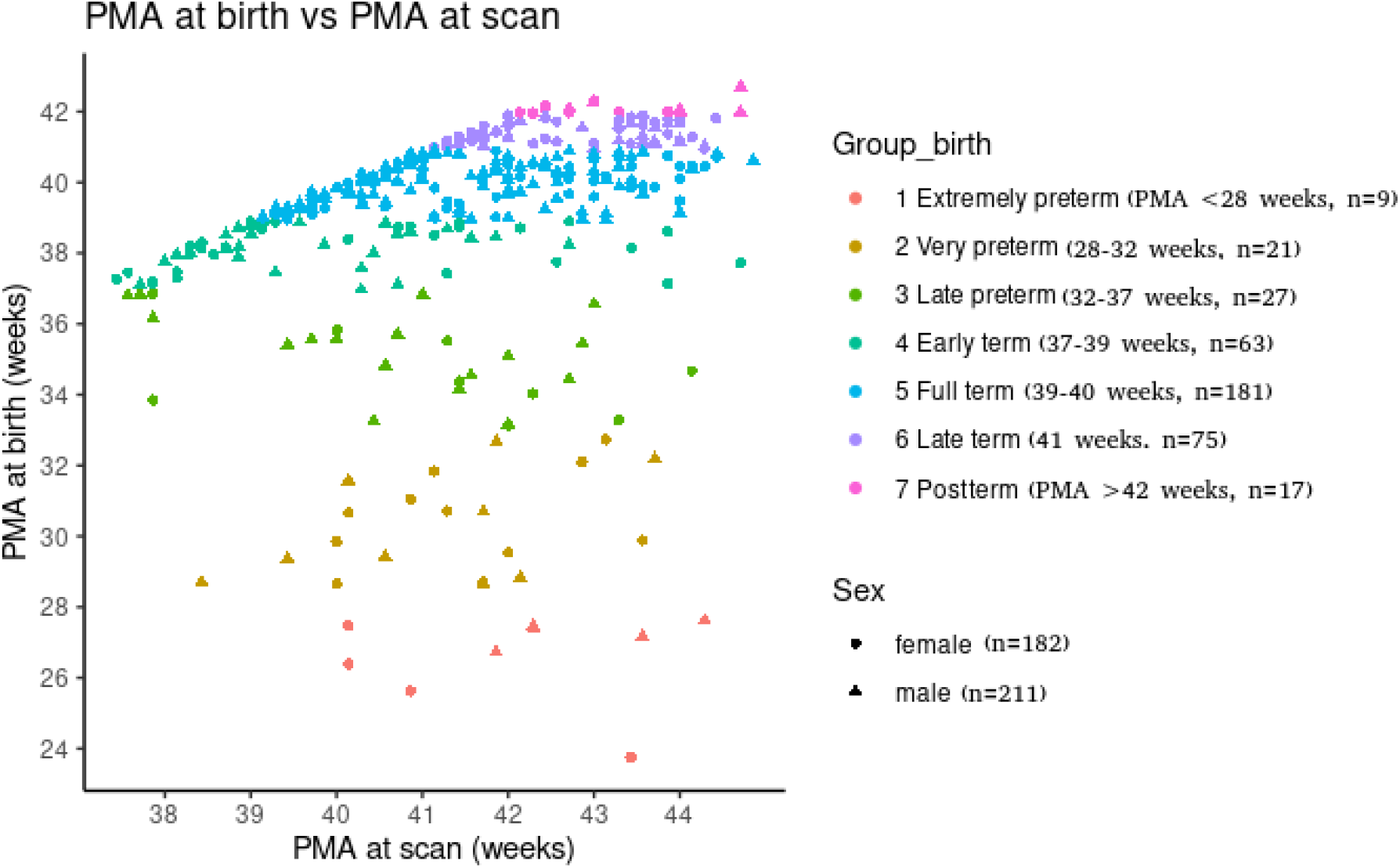
PMA at birth and at scan in cross-sectional data.

Using a one-way ANCOVA, controlling for PMA at the time of scan and sex, we compared graph theory measures among seven groups of infants as a function of their PMA at birth. A significant effect between groups was identified (Table 1, Figure 5), with post hoc tests (Tukey’s HSD) showing significant differences between preterm and term infants.

**Figure 5.**
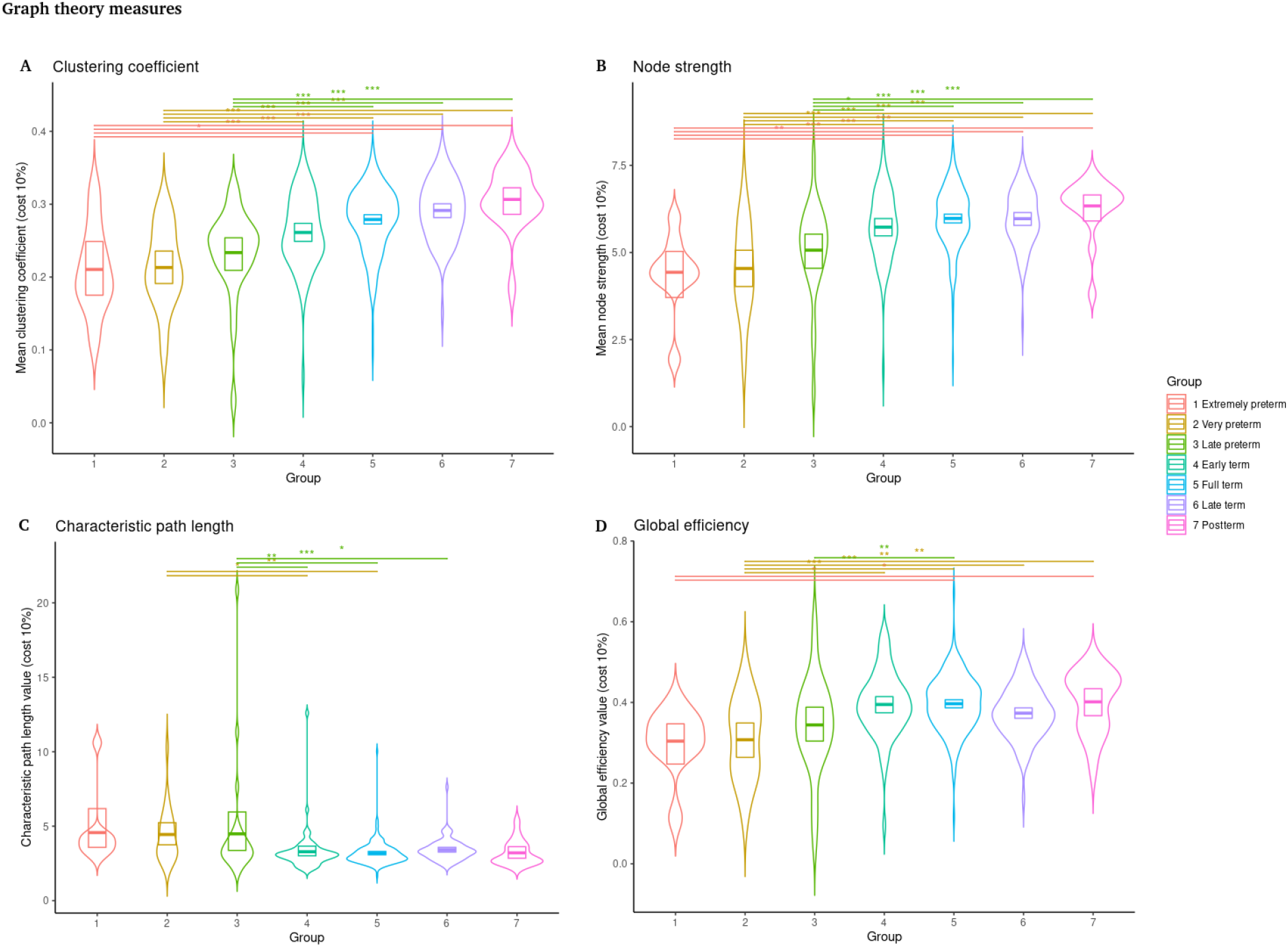
Graph theory measures. Significant differences between preterm and term infants are shown. (Lines at the top represent comparisons with p < 0.05, Tukey HSD post hoc tests). A) Mean clustering coefficient. B) Mean node strength. C) Characteristic path length. D) Global efficiency. Clustering coefficient, node strength and global efficiency increase as PMA at birth increases, controlling for age at scanning and sex, while shortest path length decreases as PMA at birth increases (* p < 0.05, ** p < 0.01, *** p < 0.001).

**Table 1.** One-way ANCOVA test for graph theory properties.

|  | Df | F value | Pr(>F) |
| --- | --- | --- | --- |
| Clustering coefficient | 6 | 16.14 | $<2 \times 10^{-16}$ *** |
| Node strength | 6 | 14.029 | $1.89 \times 10^{-14}$ *** |
| Shortest path length | 6 | 6.530 | $1.41 \times 10^{-06}$ *** |
| Global efficiency | 6 | 7.751 | $6.96 \times 10^{-08}$ *** |

The table shows the results of the 1-way ANCOVA test that compares graph theory measures between seven groups of infants based on their PMA at birth, controlling for PMA at MRI session and sex. Significant between group effect values for the four graph theory measures are shown. (\*\*\*) $p < 0.001$ .

As expected, the values for the clustering coefficient *CC*, connectivity strength *S* and global efficiency *E*_*global*_ increase as PMA at birth increases, controlling for age at scanning and sex. Additionally, the characteristic path length *d* decreases as PMA at birth increases. Figures 5A and 5B for *CC* and *S* show significant differences between preterm and term infants, while in Figures 5C and 5D *d* and *E*_*global*_ show significant differences between preterm infants and at least one group of at-term infants.

### 2. Local effects (brain regions)

In general, the results show that even at term-equivalent age, the higher the PMA at birth is associated with better global properties of the network. Then, the question if these findings are true for all the brain regions arises. Figures 6 to 8 show the graph theory parameters estimated for each brain region for the seven groups and the corresponding F value obtained by performing a one-way ANCOVA between groups, controlling for PMA at the time of scan and sex.

**Figure 6.**
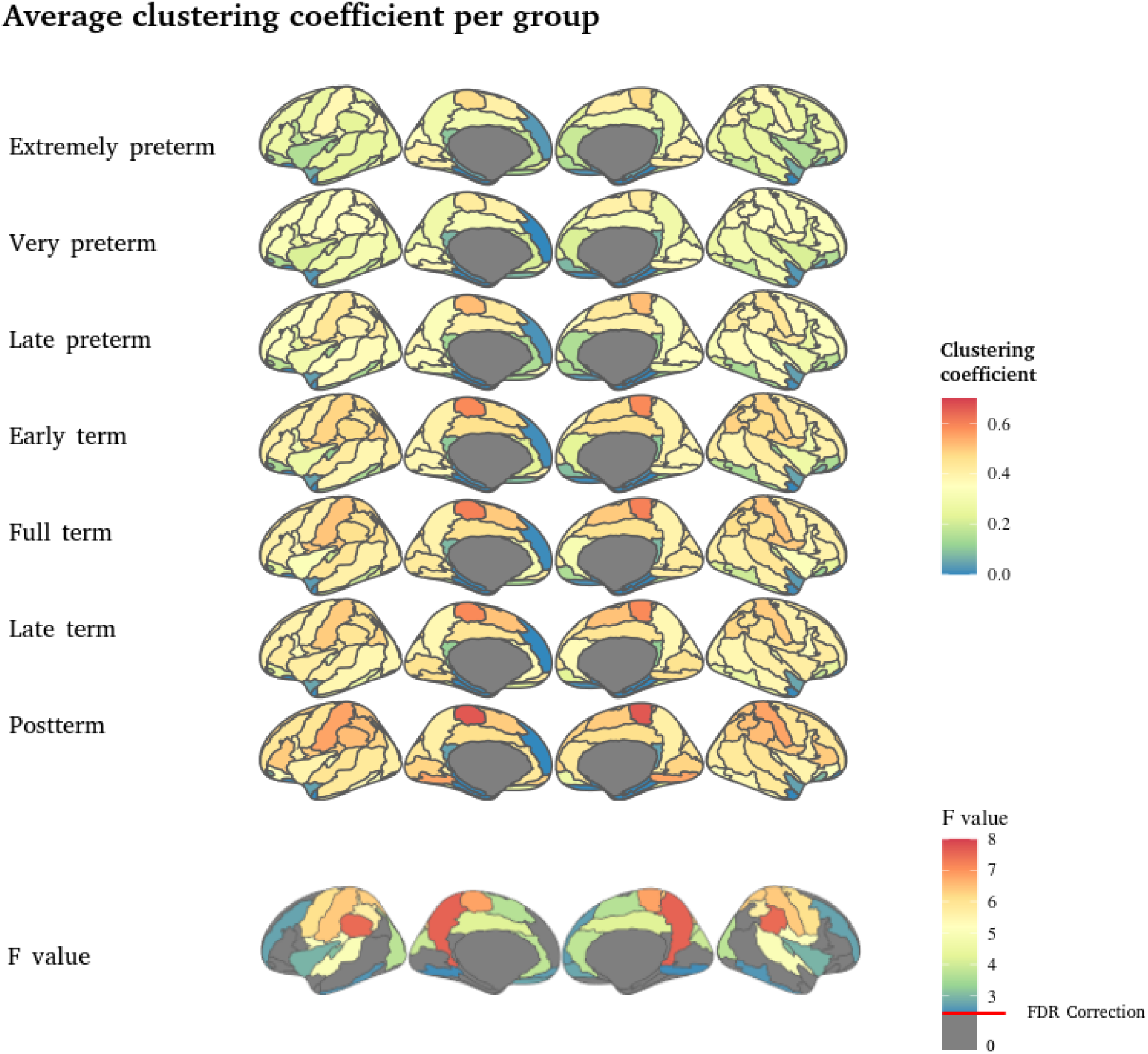
Average clustering coefficient per group. The first six rows show the average **CC** value per group; the higher the PMA, the higher the **CC** value in the central, parietal and sensorimotor areas. The seventh row shows the *F values* of the ANCOVA test between groups by brain regions; the higher the value, the more significant the difference between the groups. The red line indicates the correction with FDR at q < 0.05 (p < 0.026, F value = 2.477).

**Figure 7.**
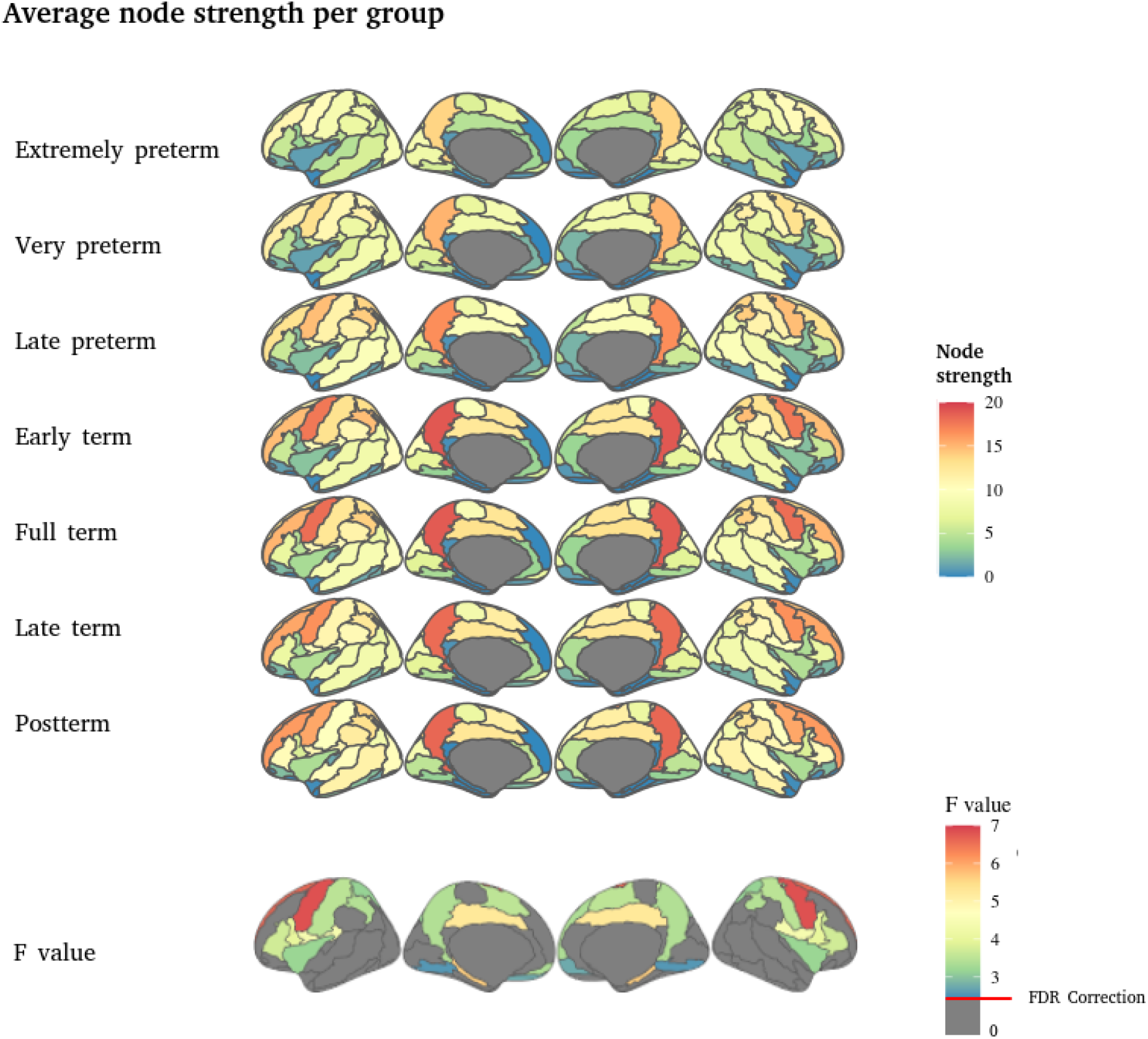
Average node strength per group. The first six rows show the mean *S* value per group; the higher the PMA, the higher the **S** value in the central areas. The seventh row shows the *F values* of the ANCOVA test between groups; the higher the value, the more significant the difference between the regions. The red line indicates the correction with FDR at q < 0.05 (p < 0.021, F value = 2.568).

**Figure 8.**
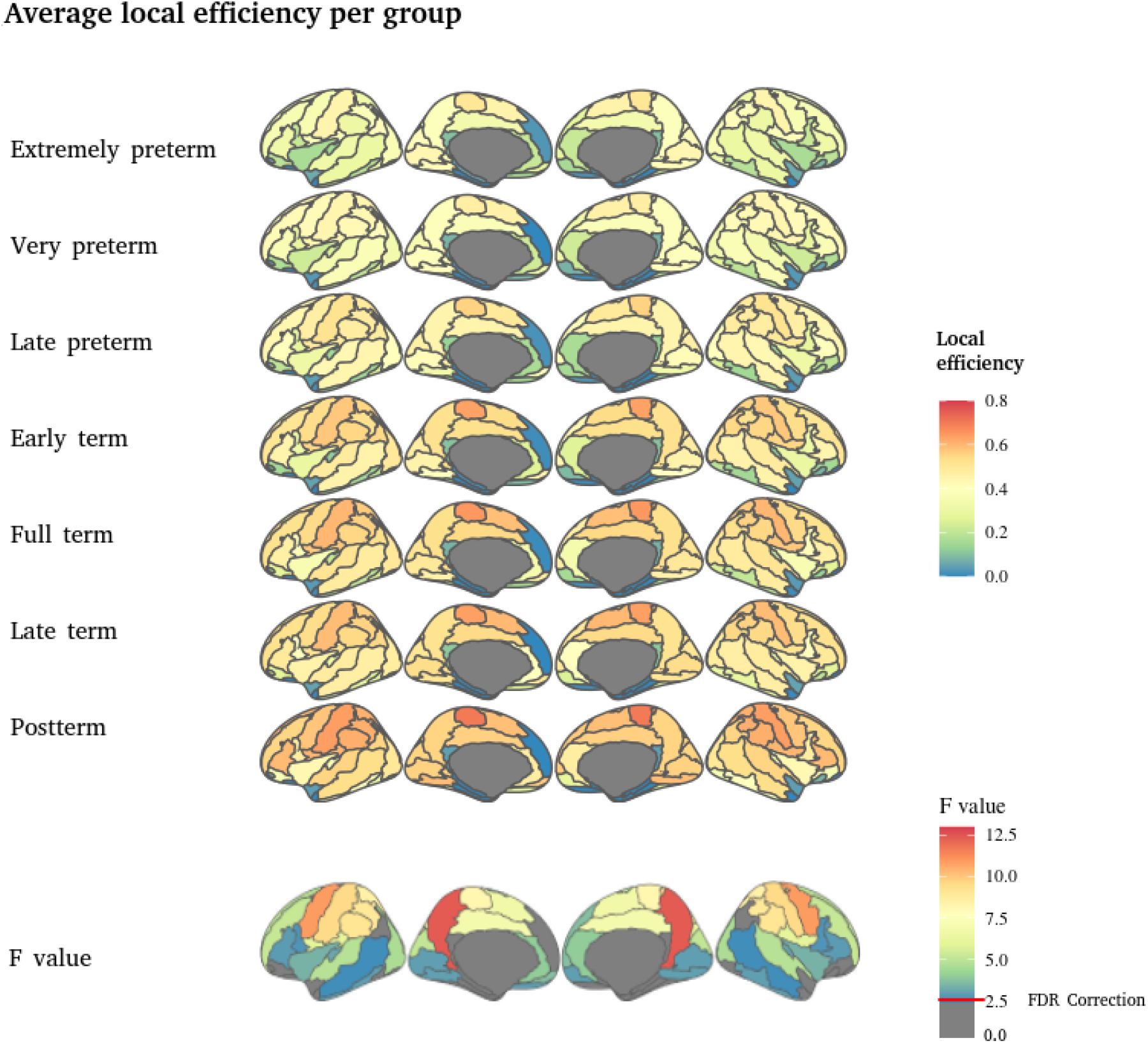
Average local efficiency per group. The first six rows show the average local efficiency **E*_*locall*_*, per group, the higher the PMA, the higher the **E*_*locall*_* value in the central and parietal areas. The seventh row shows the *F values* of the ANCOVA test between groups; the higher the value, the more significant the difference between groups. The red line indicates the correction with FDR at q < 0.05 (p < 0.03, F value = 2.534).

Figure 6 shows that the largest effects *CC* are located in the precuneus, precentral, postcentral and paracentral gyri. But a large portion of the brain show significant differences between groups, including primary sensory regions.

For the characteristic path length the most significant effects are also observed in the precuneus and the precentral and middle frontal gyri, with other regions of primary sensory regions also showing significant differences between groups (Figure 7).

The efficiency of a node was estimated on the subgraph created by its neighbors, this is defined as the local efficiency (*E*_*locall*_). Again the brain regions with the largest differences in the *E*_*locall*_ are the precuneus and the precentral, postcentral and paracentral gyri. But significant differences are identified in several brain regions, particularly including primary sensory regions (Figure 8).

### 3. Developmental trajectories

When exploring the developmental trajectories of such functional organization properties, the best fitting models showed non-linear trajectories for all the properties in preterm neonates and two of them in at-term neonates. Specifically, in at-term neonates, clustering coefficient showed quadratic fit and node strength showed a logarithmic trajectory, while the shortest path length and global efficiency showed linear trajectories (Figure 9). However, it is worth noting that all models for at-term neonates showed very similar fitting, with just marginal differences in their AIC (Table 2).

**Figure 9.**
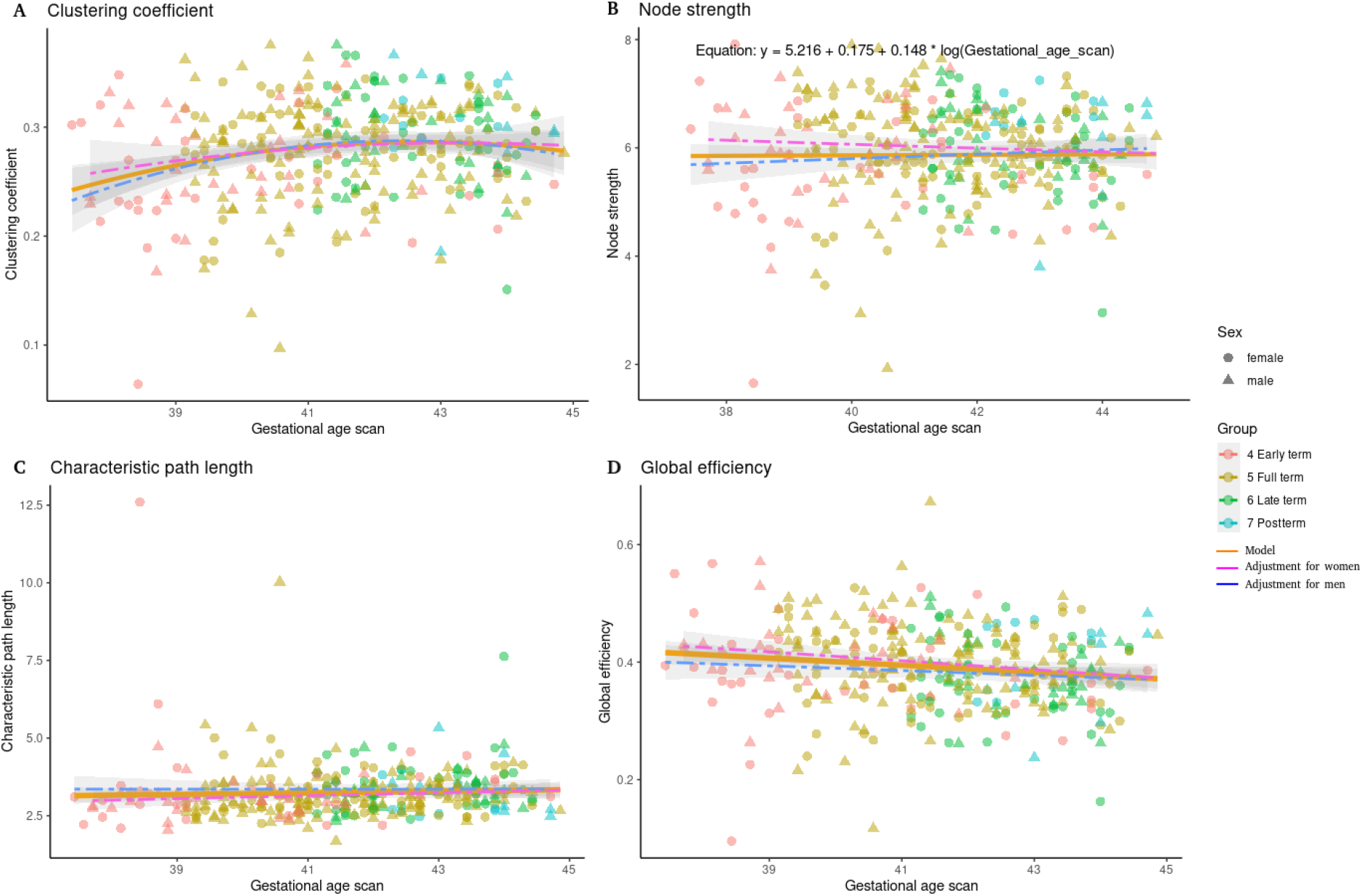
Best fitting models in at-term infants. A) Clustering coefficient. B) Node strength. C) Characteristic path length. D) Global efficiency. Clustering coefficient, node strength and global efficiency increase as gestational age scan increases.

**Figure 10.**
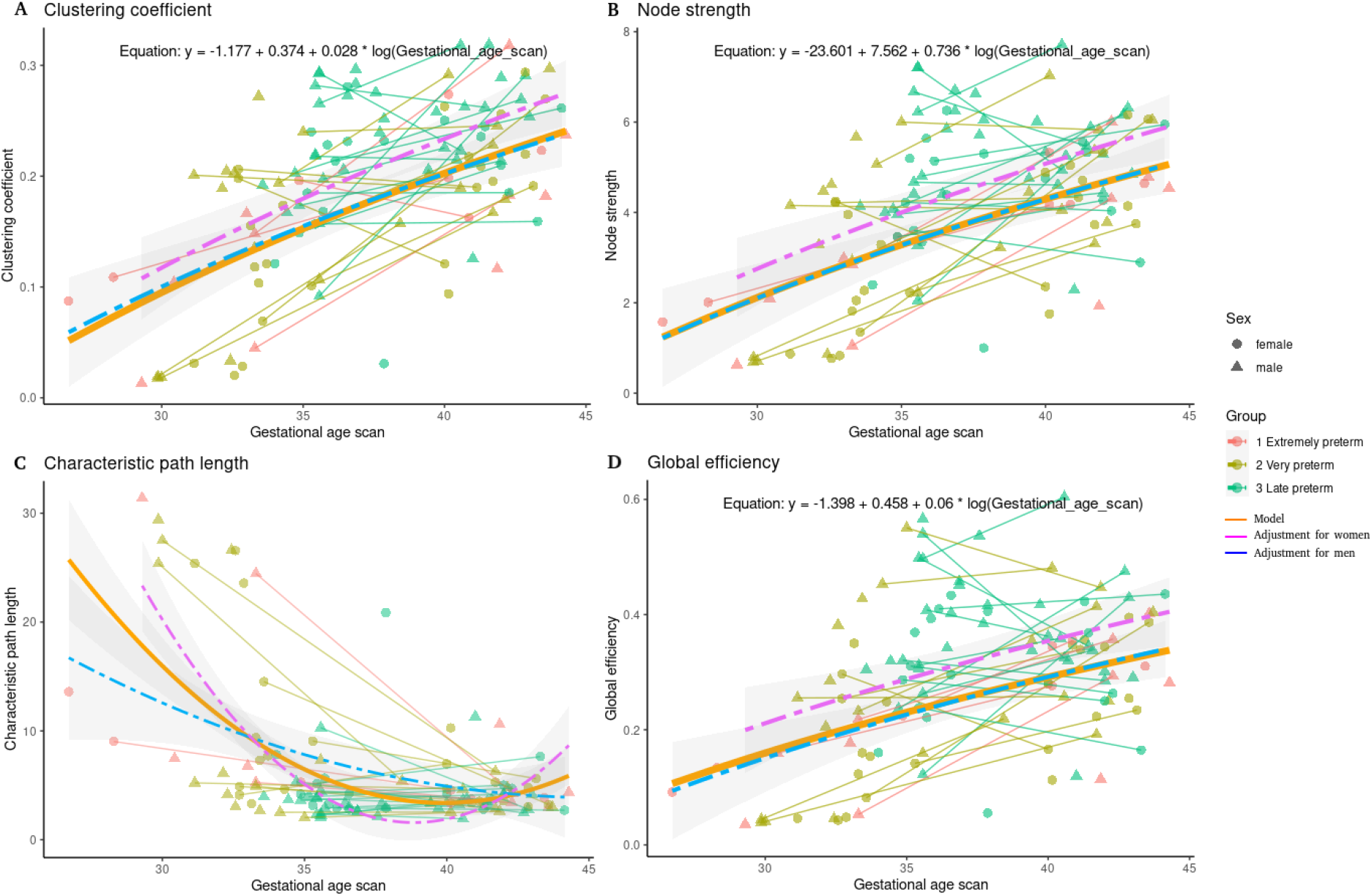
Best fitting models for the network properties in preterm infants. A) Clustering coefficient. B) Node strength. C) Characteristic path length. D) Global efficiency. Clustering coefficient, node strength and global efficiency increase as gestational age scan increases.

**Table 2.**
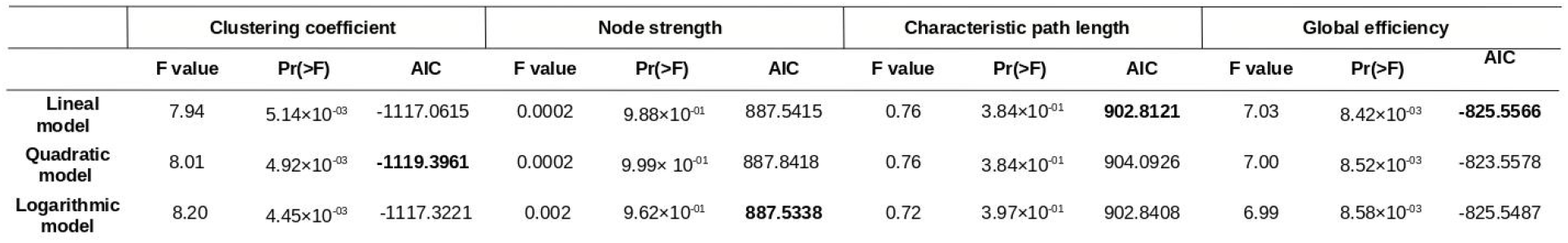
Model fitting for graph theory measures in at-term neonates.

|  | Clustering coefficient |  |  | Node strength |  |  | Characteristic path length |  |  | Global efficiency |  |  |
| --- | --- | --- | --- | --- | --- | --- | --- | --- | --- | --- | --- | --- |
|  | F value | Pr(>F) | AIC | F value | Pr(>F) | AIC | F value | Pr(>F) | AIC | F value | Pr(>F) | AIC |
| Lineal model | 7.94 | 5.14×10<sup>-03</sup> | -1117.0615 | 0.0002 | 9.88×10<sup>-01</sup> | 887.5415 | 0.76 | 3.84×10<sup>-01</sup> | **902.8121** | 7.03 | 8.42×10<sup>-03</sup> | **-825.5566** |
| Quadratic model | 8.01 | 4.92×10<sup>-03</sup> | **-1119.3961** | 0.0002 | 9.99× 10<sup>-01</sup> | 887.8418 | 0.76 | 3.84×10<sup>-01</sup> | 904.0926 | 7.00 | 8.52×10<sup>-03</sup> | -823.5578 |
| Logarithmic model | 8.20 | 4.45×10<sup>-03</sup> | -1117.3221 | 0.002 | 9.62×10<sup>-01</sup> | **887.5338** | 0.72 | 3.97×10<sup>-01</sup> | 902.8408 | 6.99 | 8.58×10<sup>-03</sup> | -825.5487 |

The table compares the linear, quadratic, and logarithmic models for each graph theory measure in at-term neonates. The lowest AIC value indicates the best fitting model, highlighted in bold for each case.

For the premature neonates, the best fitting models showed non-linear trajectories for all the explored properties. Specifically, the clustering coefficient, node strength and global efficiency showed logarithmic trajectories, while the characteristic path length showed the quadratic as the best fitting model (Figure 9; Table 3).

**Table 3.** Model fitting for graph theory measures in premature neonates.

|  | Clustering coefficient |  |  | Node strength |  |  | Characteristic path length |  |  | Global efficiency |  |  |
| --- | --- | --- | --- | --- | --- | --- | --- | --- | --- | --- | --- | --- |
|  | F value | Pr(>F) | AIC | F value | Pr(>F) | AIC | F value | Pr(>F) | AIC | F value | Pr(>F) | AIC |
| Lineal model | 57.19 | $8.92 \times 10^{-11}$ | -283.7938 | 46.14 | $2.67 \times 10^{-09}$ | 430.4784 | 35.72 | $1.15 \times 10^{-07}$ | 747.1826 | 25.99 | $2.68 \times 10^{-06}$ | -141.1299 |
| Quadratic model | 11.54 | $9.65 \times 10^{-04}$ | -275.5133 | 13.82 | $3.26 \times 10^{-04}$ | 430.5893 | 23.74 | $4.34 \times 10^{-06}$ | **735.6559** | 15.49 | $1.52 \times 10^{-04}$ | -138.0371 |
| Logarithmic model | 60.23 | $2.89 \times 10^{-11}$ | **-293.1881** | 49.06 | $9.08 \times 10^{-10}$ | **421.0729** | 39.52 | $2.97 \times 10^{-08}$ | 737.1098 | 28.28 | $1.07 \times 10^{-06}$ | **-150.2163** |

The table compares the linear, quadratic, and logarithmic fit models for each graph theory measure in preterm. The lowest AIC value indicates the best fitting model; it is highlighted in bold.

When compared by sex, at-term infants showed no significant differences between males and females, while female preterm infants showed increased clustering coefficient (p < 0.02), node strength (p < 0.01) and global efficiency (p < 0.01). We observed a potential interaction between sex and age in the graphs, so we tested these additional models but found no significant interaction.

### 4. Longitudinal dataset

Subsequently, we wanted to investigate how global network characteristics change with age in the subjects with more than one acquisition. For this, we used the longitudinal subsample of 37 infants with two MRI scans each and without radiological signs of white matter lesions (Figure 11).

**Figure 11.**
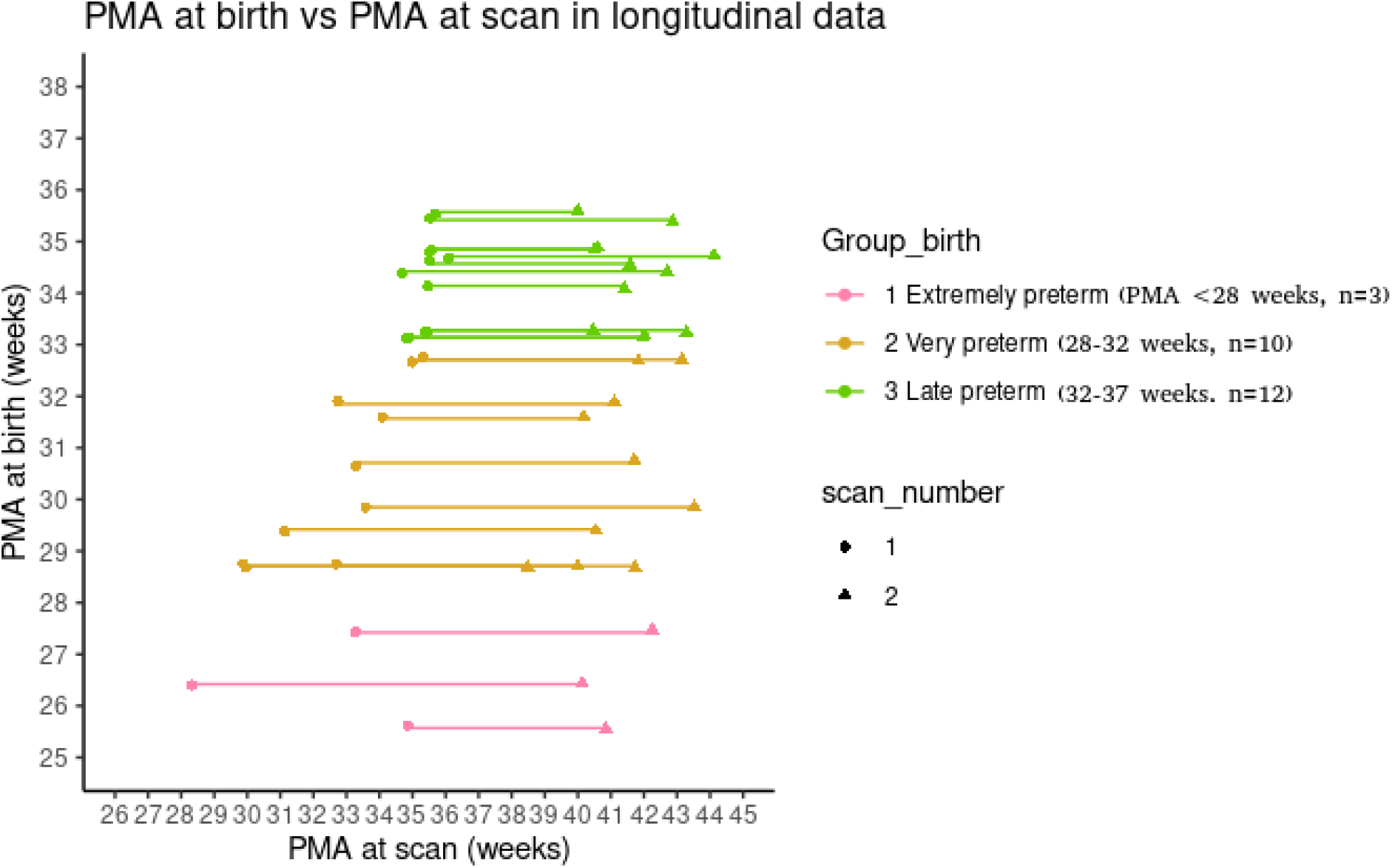
PMA at birth and at scan for the longitudinal data. Lines represent longitudinal data for the same participant.

The following results are also shown with a connection density of 10%. Using a Wilcoxon test for repeated measures, we compared the values of the graph theory measures between the first and second scan in the extremely preterm, very preterm and late preterm groups.

There is a tendency to improve values in all measures of graph theory in the second scan (Figure 12); however, there is noticeable variability among preterm infants, with some infants showing no changes with time or even the opposite change to that of the group. The number of extremely preterm infants is very small and this may limit the finding of significant differences due to small statistical power; however, the second imaging session tended to have improved values (at term-equivalent age) for all the measures. On the other hand, the sample of late preterm infants only shows significant differences for clustering coefficient, potentially due to the few days between scanning sessions. Figure 12 shows the results for the four measurements, the figure 12A is for *CC*, no significant differences are observed in extremely preterms t = -1.2746, df = 2, p-value = 0.3305, while in very preterm t = -2.4095, df = 9, p-value = 0.03928 and late preterm infants t = -2.2194, df = 11, p-value = 0.04842 there are significant differences. As expected, *CC* increases with increasing PMA, the network seems to be more segregated in the second resonance at term-equivalent age.

**Figure 12.**
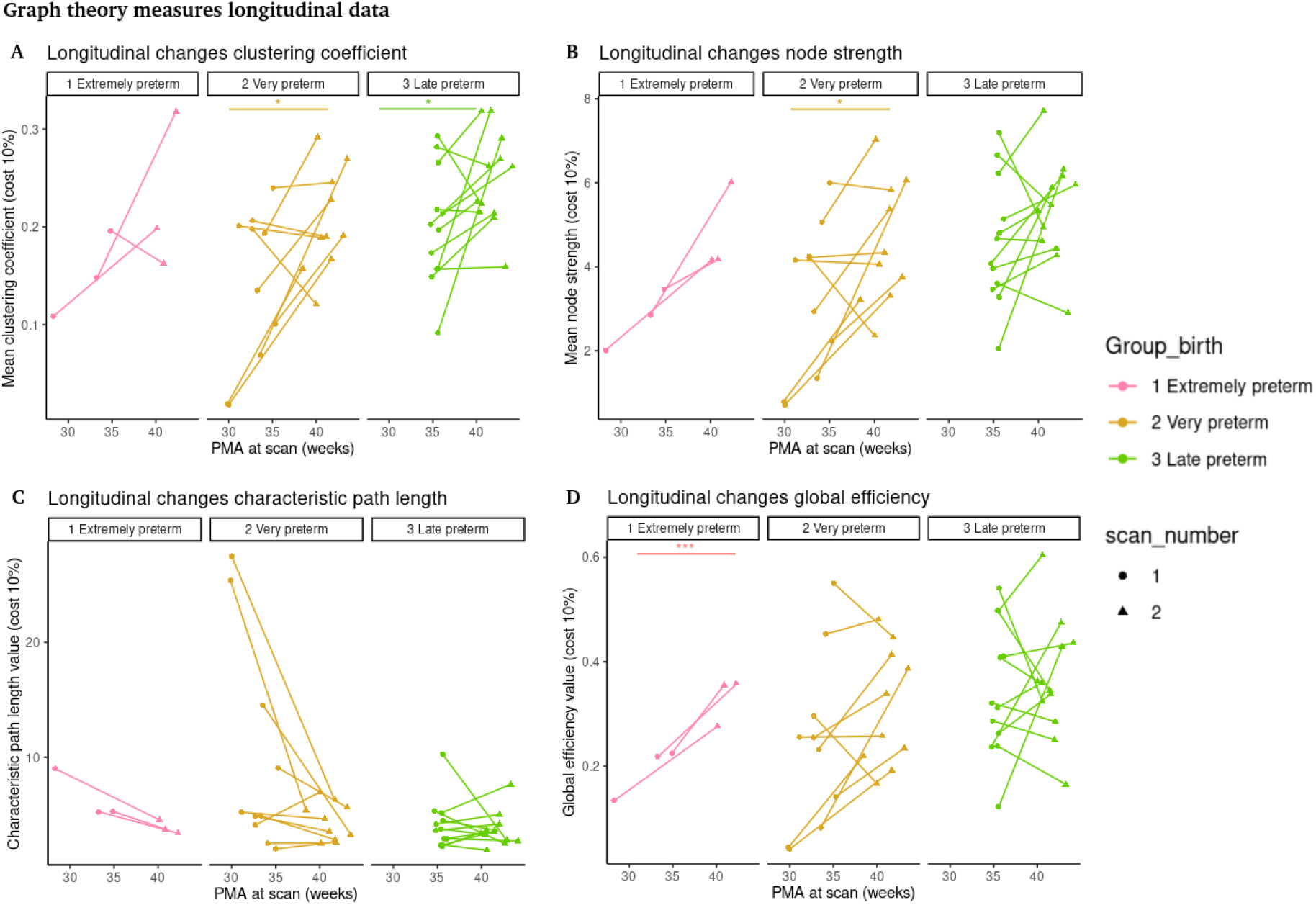
Longitudinal data for graph theory measures. A) Longitudinal changes in clustering coefficient. B) Longitudinal changes in node strength. C) Longitudinal changes in characteristic path length. D) Longitudinal changes in global efficiency. Lines and the star at the top represent comparisons with p < 0.05.

Similarly, the connectivity strength also shows significant differences only for very preterm infants (extremely preterm t = -2.8229, df = 2, p-value = 0.1059; very preterm t= -2.2859, df = 9, p-value = 0.0481; late preterm t = -1.4559, df = 11, p-value = 0.1734; Figure 12B). The same is true for the characteristic path length *d* (extremely preterm t = =-2.8229, df = 2, p-value = 0.1059; very preterm t = 2.047, df = 9, p-value = 0.07095; late preterm t = 0.60867, df = 11, p-value = 0.5551; Figure 12C).

Finally, for the global efficiency *E*_*global*_, no significant differences were observed for any group (extremely preterm t = -39.064, df = 2, p-value = 0.0006547; very preterm t = -1.8521, df = 9, p-value = 0.09703; late preterm t = -0.46267, df = 11, p-value = 0.6526; Figure 12D).

## Discussion

In this study, we performed a detailed analysis of the functional organization of the brain and its developmental trajectory in preterm and at-term neonates.

We performed a detailed exploration on the effects of PMA at birth, first dividing the sample in a total of seven groups: three preterm groups categorized by the WHO criteria and four at-term groups categorized according to the guidelines of the American College of Obstetricians and Gynecologists and the Society for Maternal-Fetal Medicine. The results show significant and robust differences for the premature neonates even at term-equivalent age, with an evident gradient of effects depending on the PMA at birth. The results are reproduced for a large variety of connectivity thresholds, here density thresholds ranging from the top 5% higher connectivity values to the 50% with 5% intervals, evidence that these results are very robust independently of the connectivity threshold.

Based on the classification of the American College of Obstetricians and Gynecologists and the Society for Maternal-Fetal Medicine, there has been a debate whether the early term group presents sufficient fetal maturity to avoid the neurological comorbidities of prematurity. Recent research on this specific group of early term subjects has revealed risk factors for physiological comorbidities, such as respiratory deficiency and diabetes that seem to persist into adulthood (Chen et al., 2022; Odibo et al., 2016). However, our results on the brain functional organization suggest that this group is more similar to the other at-term groups rather than to the premature groups. Specifically, the early terms showed significant differences with at least one of the preterm groups in all the properties here explored, but showed no significant difference with any of the other three at-term groups.

When exploring such properties at the local level (brain regions), the precuneus, mid cingulate, primary motor and somatosensory cortices, parietal gyrus, orbitofrontal cortex, superior and inferior frontal gyrus, temporal gyrus, occipital gyrus, supramarginal gyrus, Heschl gyrus left, Rolandic operculum, lingual gyrus, cingulate gyrus and insula showed the greatest differences between groups (Supplementary Material). It is not surprising that these regions include the primary somatosensory and primary motor regions, as premature birth is usually associated with higher risks for sensorimotor, auditory, visual, cognitive and language deficits, as well as for ADHD, ASD and CP (Chen et al., 2022; Do et al., 2019; Hee Chung et al., 2020). As such, most of the prospective interventions in preterm infants include sensorimotor enrichment (Harmony et al., 2016), with overall benefits in behavioral and cognitive domains. In particular, the auditory cortex showed the greatest effect in the left hemisphere, which is largely recognized as the commonly dominant hemisphere for language acquisition (Gracia-Tabuenca et al., 2018; Olulade et al., 2020). This result provides the potential neurofunctional substrate for the language acquisition deficits identified in preterm infants (Gozzo et al., 2009; Myers et al., 2010; Salvan et al., 2017; Vandormael et al., 2019). In addition, the largest differences between groups were identified for the precuneus (Figures 6-8), which is one of the main hubs of the brain network and part of the DMN. Alterations in the connectivity of this network have been associated with ADHD (Castellanos & Aoki, 2016; Elton et al., 2014; Gracia-Tabuenca et al., 2020), visuospatial abilities (Fernandez-Baizan et al., 2021; Woodward et al., 2021) episodic memory (Schommartz et al., n.d.; Stedall et al., 2023), affective disorders (Guilherme Monte Cassiano et al., 2016) and social cognition (Pereira et al., 2017; You et al., 2019). Our results suggest a link between these brain functional alterations and the vulnerability to develop such behavioral disorders.

To characterize the developmental trajectories of the graph theory properties explored here, linear, quadratic, and logarithmic models were fitted using gestational age at the scanning session as a fixed effect, with subject ID as a random variable and sex as confounding variable. The best fitting models for preterms were logarithmic, except for the characteristic path length that followed a quadratic trajectory (Table 3). Sex showed significant effect in clustering coefficient, node strength and global efficiency, showing greater values for females. This evidence seems to be consistent with recent work noting that females at term-equivalent age show increased connectivity of inferior occipitotemporal regions within the visual association network (Eyre et al., 2021). In at-term neonates, clustering coefficient showed quadratic fit and node strength showed a logarithmic trajectory, while the characteristic path length and global efficiency showed linear trajectories. However, we must note that for the at-term neonates the AIC was very similar for all the tested models for each property (Table 2). These results are relevant as we identified no previous description of such trajectories for these properties, with only one recent study suggesting non-linear trajectories for the development of resting state networks in the same sample (Kim et al., 2023).

Finally, using the longitudinal subsample of preterm infants, we identified a significant increase with age from ages lower than 37 WG to at term-equivalent ages, mainly for the very preterm group in all properties, except for the global efficiency (Figure 12). Global efficiency is considered a measure of network integration, which in the brain is understood to be supported by the establishment of long-range connections (Cao et al., 2016; Gao et al., 2011). The emergence of these connections has been associated with the development of the white matter microstructural properties, including myelination (Dubois et al., 2014). However, premature birth is known to alter the production of premyelinating oligodendrocytes during gestation, which consequently alters myelin production in later stages of development (Volpe, 2009). This interruption in myelin maturation may be associated with the lack of significant changes in the global efficiency of the network (Figure 12). In contrast, extremely preterm infants showed no significant changes, potentially due to the small sample size for this group. Meanwhile, the late preterm infants showed significant changes only for the clustering coefficient, probably due to the shortest period between both scans, preventing the observation of more notable changes. In general, a high degree of variability in the developmental changes can be observed in this longitudinal subsample, with some of the subjects showing no change or even opposing effects to the group average change. This variability may be associated with the varying prevalence rates for a diversity of developmental outcomes. However, results from cognitive-behavioral assessments and demographic data were not available to explore these associations.

Overall, our results confirm that the functional connectivity, the integration and segregation properties of the premature brain are significantly decreased at term-equivalent age, despite showing rapid maturation after birth.

## Conclusions

Preterm birth is associated with significant differences in the functional organization of the brain network compared with at-term neonates, even in the absence of evident brain injuries. Dividing the sample into finer groups allowed us to observe with greater detail how the smaller the PMA, the greater the differences in the functional network even at term-equivalent ages. However, the premature infants show dramatic improvement in the brain network from birth to at term-equivalent age with a clear sexual dimorphism and important variability among the individual developmental trajectories.

## Acknowledgments

Nelsiyamid López-Guerrero is a doctoral student from the Programa de Doctorado en Ciencias Biomédicas, at the Universidad Nacional Autónoma de México (UNAM) and received the fellowship 823584 from CONAHCYT, Mexico. This work was supported by the grant UNAM-PAPIIT IN208622 to Sarael Alcauter. We thank Leopoldo González Santos and the Unidad de Cómputo del Instituto de Neurobiología, UNAM, for their technical assistance and Michael C. Jeziorski for editing this manuscript. Data was provided by the developing Human Connectome Project, KCL-Imperial-Oxford Consortium funded by the European Research Council under the European Union Seventh Framework Programme (FP/2007-2013) / ERC Grant Agreement no. [319456]. We are grateful to the families who generously supported this trial and the dHCP for sharing the datasets.

