## Supplementary Material for "Developmental trajectories and differences in functional brain network properties of preterm and at-term neonates"

**Cross-sectional age term-equivalent dataset**

Table S1. Premature infants

| ID | sesion |
|---|---|
| sub-CC00271XX08 | 100400 |
| sub-CC00284AN13 | 111400 |
| sub-CC00287AN16 | 91800 |
| sub-CC00287BN16 | 93900 |
| sub-CC00290XX11 | 92900 |
| sub-CC00301XX04 | 113001 |
| sub-CC00305XX08 | 115700 |
| sub-CC00309BN12 | 99400 |
| sub-CC00326XX13 | 118800 |
| sub-CC00385XX15 | 125700 |
| sub-CC00388XX18 | 118700 |
| sub-CC00389XX19 | 133800 |
| sub-CC00407BN11 | 137200 |
| sub-CC00465XX12 | 137900 |
| sub-CC00525XX14 | 165900 |
| sub-CC00529AN18 | 170000 |
| sub-CC00529BN18 | 170100 |
| sub-CC00569XX17 | 170600 |
| sub-CC00572BN12 | 161400 |
| sub-CC00576XX16 | 178200 |
| sub-CC00617XX15 | 188400 |
| sub-CC00618XX16 | 193600 |
| sub-CC00628XX18 | 193500 |
| sub-CC00630XX12 | 182500 |
| sub-CC00632XX14 | 196000 |
| sub-CC00648XX22 | 204400 |
| sub-CC00672AN13 | 214900 |
| sub-CC00672BN13 | 214800 |
| sub-CC00712XX11 | 232701 |
| sub-CC00747XX22 | 238600 |
| sub-CC00768XX18 | 236400 |
| sub-CC00770XX12 | 1100 |
| sub-CC00771XX13 | 17710 |
| sub-CC00792XX18 | 1800 |
| sub-CC00802XX10 | 11210 |
| sub-CC00823XX15 | 27810 |
| sub-CC00845AN21 | 32010 |
| sub-CC00845BN21 | 32110 |
| sub-CC00855XX14 | 530 |
| sub-CC00867XX18 | 8930 |
| sub-CC00889AN24 | 9230 |
| sub-CC00889BN24 | 9330 |
| sub-CC00891XX18 | 30630 |
| sub-CC00907XX16 | 19230 |

| | |
|---|---|
| sub-CC00919XX20 | 23630 |
| sub-CC00945AN22 | 13730 |
| sub-CC00997BN25 | 56430 |
| sub-CC01005XX07 | 49930 |
| sub-CC01011XX05 | 55231 |
| sub-CC01038XX16 | 67330 |
| sub-CC01059AN12 | 72130 |
| sub-CC01059CN12 | 72230 |
| sub-CC01077XX14 | 78430 |
| sub-CC01104XX07 | 82830 |
| sub-CC01111XX06 | 100031 |
| sub-CC01209XX13 | 144830 |
| sub-CC01218XX14 | 157231 |

Table S2. At-term infants

| ID | sesion |
|---|---|
| sub-CC00257XX10 | 84700 |
| sub-CC00258XX11 | 84900 |
| sub-CC00260XX05 | 85300 |
| sub-CC00261XX06 | 85600 |
| sub-CC00264AN09 | 89400 |
| sub-CC00265XX10 | 86901 |
| sub-CC00267XX12 | 87700 |
| sub-CC00268XX13 | 87800 |
| sub-CC00269XX14 | 88300 |
| sub-CC00272XX09 | 117900 |
| sub-CC00275XX12 | 93501 |
| sub-CC00286XX15 | 91700 |
| sub-CC00289XX18 | 119700 |
| sub-CC00295XX16 | 127500 |
| sub-CC00298XX19 | 94700 |
| sub-CC00299XX20 | 94900 |
| sub-CC00300XX03 | 96000 |
| sub-CC00302XX05 | 113500 |
| sub-CC00303XX06 | 96900 |
| sub-CC00304XX07 | 111600 |
| sub-CC00306XX09 | 98700 |
| sub-CC00307XX10 | 98800 |
| sub-CC00308XX11 | 98900 |
| sub-CC00313XX08 | 100000 |
| sub-CC00314XX09 | 100101 |
| sub-CC00316XX11 | 101300 |
| sub-CC00319XX14 | 117300 |
| sub-CC00320XX07 | 102300 |
| sub-CC00324XX11 | 111200 |
| sub-CC00328XX15 | 104800 |
| sub-CC00330XX09 | 128200 |
| sub-CC00332XX11 | 105700 |
| sub-CC00334XX13 | 106100 |
| sub-CC00335XX14 | 106300 |
| sub-CC00336XX15 | 106600 |
| sub-CC00337XX16 | 107000 |
| sub-CC00338AN17 | 107700 |

| | |
|---|---|
| sub-CC00338BN17 | 107600 |
| sub-CC00339XX18 | 107200 |
| sub-CC00340XX11 | 107902 |
| sub-CC00341XX12 | 108000 |
| sub-CC00343XX14 | 108500 |
| sub-CC00344XX15 | 108600 |
| sub-CC00345XX16 | 109000 |
| sub-CC00347XX18 | 109600 |
| sub-CC00348XX19 | 110200 |
| sub-CC00349XX20 | 110300 |
| sub-CC00352XX06 | 110700 |
| sub-CC00354XX08 | 112100 |
| sub-CC00355XX09 | 112600 |
| sub-CC00356XX10 | 112901 |
| sub-CC00357XX11 | 113900 |
| sub-CC00358XX12 | 114400 |
| sub-CC00362XX08 | 114500 |
| sub-CC00363XX09 | 114900 |
| sub-CC00364XX10 | 115200 |
| sub-CC00367XX13 | 116000 |
| sub-CC00368XX14 | 116600 |
| sub-CC00370XX08 | 142400 |
| sub-CC00371XX09 | 134700 |
| sub-CC00376XX14 | 118400 |
| sub-CC00377XX15 | 119800 |
| sub-CC00378XX16 | 120200 |
| sub-CC00380XX10 | 121200 |
| sub-CC00381XX11 | 121600 |
| sub-CC00382XX12 | 121700 |
| sub-CC00383XX13 | 121800 |
| sub-CC00400XX04 | 123700 |
| sub-CC00401XX05 | 123900 |
| sub-CC00402XX06 | 124300 |
| sub-CC00403XX07 | 124400 |
| sub-CC00404XX08 | 124500 |
| sub-CC00408XX12 | 125500 |
| sub-CC00409XX13 | 125600 |
| sub-CC00410XX06 | 125800 |
| sub-CC00412XX08 | 126301 |
| sub-CC00413XX09 | 127100 |
| sub-CC00415XX11 | 127400 |
| sub-CC00421AN09 | 126000 |
| sub-CC00424XX12 | 129400 |
| sub-CC00425XX13 | 129800 |
| sub-CC00426XX14 | 129900 |
| sub-CC00427XX15 | 130100 |
| sub-CC00428XX16 | 130400 |
| sub-CC00429XX17 | 130900 |
| sub-CC00430XX10 | 131100 |
| sub-CC00431XX11 | 131500 |
| sub-CC00433XX13 | 132000 |
| sub-CC00438XX18 | 156800 |

| | |
|---|---|
| sub-CC00440XX12 | 132200 |
| sub-CC00441XX13 | 132902 |
| sub-CC00442XX14 | 133300 |
| sub-CC00443XX15 | 133900 |
| sub-CC00444XX16 | 135101 |
| sub-CC00445XX17 | 134800 |
| sub-CC00446XX18 | 135200 |
| sub-CC00447XX19 | 135600 |
| sub-CC00448XX20 | 135800 |
| sub-CC00450XX05 | 136100 |
| sub-CC00451XX06 | 137000 |
| sub-CC00453XX08 | 136600 |
| sub-CC00455XX10 | 137700 |
| sub-CC00457XX12 | 138601 |
| sub-CC00458XX13 | 138900 |
| sub-CC00461XX08 | 175100 |
| sub-CC00466AN13 | 138400 |
| sub-CC00466BN13 | 138300 |
| sub-CC00467XX14 | 139000 |
| sub-CC00468XX15 | 139100 |
| sub-CC00469XX16 | 139200 |
| sub-CC00470XX09 | 139600 |
| sub-CC00472XX11 | 140000 |
| sub-CC00473XX12 | 140100 |
| sub-CC00474XX13 | 140500 |
| sub-CC00475XX14 | 141400 |
| sub-CC00476XX15 | 141500 |
| sub-CC00477XX16 | 141600 |
| sub-CC00478XX17 | 141601 |
| sub-CC00479XX18 | 141608 |
| sub-CC00480XX11 | 141609 |
| sub-CC00481XX12 | 141800 |
| sub-CC00482XX13 | 142000 |
| sub-CC00483XX14 | 144200 |
| sub-CC00484XX15 | 142700 |
| sub-CC00485XX16 | 143100 |
| sub-CC00486XX17 | 144300 |
| sub-CC00497XX20 | 144500 |
| sub-CC00498XX21 | 144900 |
| sub-CC00499XX22 | 145800 |
| sub-CC00500XX05 | 145900 |
| sub-CC00501XX06 | 146500 |
| sub-CC00502XX07 | 146700 |
| sub-CC00504XX09 | 146800 |
| sub-CC00505XX10 | 146900 |
| sub-CC00506XX11 | 147600 |
| sub-CC00507XX12 | 148202 |
| sub-CC00508XX13 | 148700 |
| sub-CC00509XX14 | 148800 |
| sub-CC00512XX09 | 150700 |
| sub-CC00514XX11 | 151400 |
| sub-CC00515XX12 | 152600 |

| | |
|---|---|
| sub-CC00516XX13 | 152902 |
| sub-CC00520XX09 | 150201 |
| sub-CC00528XX17 | 183200 |
| sub-CC00532XX13 | 154200 |
| sub-CC00534XX15 | 155100 |
| sub-CC00535XX16 | 155600 |
| sub-CC00536XX17 | 156900 |
| sub-CC00537XX18 | 157100 |
| sub-CC00538XX19 | 157200 |
| sub-CC00541XX14 | 165500 |
| sub-CC00542XX15 | 165800 |
| sub-CC00544XX17 | 169300 |
| sub-CC00548XX21 | 157400 |
| sub-CC00549XX22 | 157600 |
| sub-CC00550XX06 | 157800 |
| sub-CC00551XX07 | 158001 |
| sub-CC00552XX08 | 159300 |
| sub-CC00553XX09 | 159000 |
| sub-CC00554XX10 | 160200 |
| sub-CC00557XX13 | 163400 |
| sub-CC00560XX08 | 159800 |
| sub-CC00561XX09 | 159900 |
| sub-CC00564XX12 | 154100 |
| sub-CC00566XX14 | 164500 |
| sub-CC00568XX16 | 198900 |
| sub-CC00577XX17 | 180400 |
| sub-CC00580XX12 | 173700 |
| sub-CC00581XX13 | 177101 |
| sub-CC00583XX15 | 178600 |
| sub-CC00584XX16 | 178800 |
| sub-CC00586XX18 | 179000 |
| sub-CC00588XX20 | 183600 |
| sub-CC00589XX21 | 184000 |
| sub-CC00590XX14 | 187000 |
| sub-CC00592XX16 | 188800 |
| sub-CC00593XX17 | 189401 |
| sub-CC00594XX18 | 189800 |
| sub-CC00596XX20 | 190100 |
| sub-CC00597XX21 | 190200 |
| sub-CC00598XX22 | 190300 |
| sub-CC00607XX13 | 179300 |
| sub-CC00616XX14 | 184700 |
| sub-CC00620XX10 | 210900 |
| sub-CC00622XX12 | 185600 |
| sub-CC00637XX19 | 195700 |
| sub-CC00639XX21 | 216800 |
| sub-CC00642XX16 | 218100 |
| sub-CC00647XX21 | 190801 |
| sub-CC00649XX23 | 191201 |
| sub-CC00650XX07 | 218007 |
| sub-CC00652XX09 | 191600 |
| sub-CC00653XX10 | 191801 |

| | |
|---|---|
| sub-CC00654XX11 | 192000 |
| sub-CC00655XX12 | 215600 |
| sub-CC00656XX13 | 217601 |
| sub-CC00663XX12 | 195000 |
| sub-CC00664XX13 | 195200 |
| sub-CC00667XX16 | 196800 |
| sub-CC00668XX17 | 220700 |
| sub-CC00669XX18 | 214300 |
| sub-CC00671XX12 | 197400 |
| sub-CC00685XX18 | 226100 |
| sub-CC00687XX20 | 199000 |
| sub-CC00692XX17 | 200301 |
| sub-CC00693XX18 | 201000 |
| sub-CC00698XX23 | 220400 |
| sub-CC00705XX12 | 226000 |
| sub-CC00713XX12 | 229000 |
| sub-CC00714XX13 | 240900 |
| sub-CC00716XX15 | 222800 |
| sub-CC00719XX18 | 210600 |
| sub-CC00720XX11 | 211101 |
| sub-CC00731XX14 | 214500 |
| sub-CC00734XX17 | 216900 |
| sub-CC00736XX19 | 247000 |
| sub-CC00737XX20 | 244300 |
| sub-CC00740XX15 | 238400 |
| sub-CC00741XX16 | 218900 |
| sub-CC00749XX24 | 2600 |
| sub-CC00753XX11 | 223600 |
| sub-CC00757XX15 | 12010 |
| sub-CC00765XX15 | 8210 |
| sub-CC00769XX19 | 4400 |
| sub-CC00777XX19 | 239102 |
| sub-CC00782XX16 | 240100 |
| sub-CC00783XX17 | 15910 |
| sub-CC00789XX23 | 21110 |
| sub-CC00791XX17 | 27611 |
| sub-CC00798XX24 | 245400 |
| sub-CC00799XX25 | 23810 |
| sub-CC00801XX09 | 29510 |
| sub-CC00810XX10 | 29010 |
| sub-CC00815XX15 | 4120 |
| sub-CC00816XX16 | 40010 |
| sub-CC00818XX18 | 4020 |
| sub-CC00822XX14 | 15710 |
| sub-CC00824XX16 | 38310 |
| sub-CC00839XX23 | 23710 |
| sub-CC00840XX16 | 24910 |
| sub-CC00841XX17 | 1730 |
| sub-CC00843XX19 | 4330 |
| sub-CC00846XX22 | 26710 |
| sub-CC00847XX23 | 26910 |
| sub-CC00850XX09 | 4930 |

| | |
|---|---|
| sub-CC00851XX10 | 42010 |
| sub-CC00852XX11 | 28210 |
| sub-CC00854XX13 | 41810 |
| sub-CC00856XX15 | 15330 |
| sub-CC00858XX17 | 32210 |
| sub-CC00860XX11 | 7030 |
| sub-CC00861XX12 | 20830 |
| sub-CC00863XX14 | 34810 |
| sub-CC00868XX19 | 16530 |
| sub-CC00871XX14 | 38810 |
| sub-CC00874XX17 | 18630 |
| sub-CC00875XX18 | 16630 |
| sub-CC00879XX22 | 7430 |
| sub-CC00881XX16 | 20230 |
| sub-CC00882XX17 | 13030 |
| sub-CC00884XX19 | 17230 |
| sub-CC00890XX17 | 13330 |
| sub-CC00897XX24 | 32830 |
| sub-CC00898XX25 | 33330 |
| sub-CC00908XX17 | 37030 |
| sub-CC00911XX12 | 18830 |
| sub-CC00914XX15 | 30130 |
| sub-CC00915XX16 | 18530 |
| sub-CC00917XX18 | 39130 |
| sub-CC00923XX16 | 48430 |
| sub-CC00924XX17 | 36330 |
| sub-CC00925XX18 | 46230 |
| sub-CC00928XX21 | 37430 |
| sub-CC00929XX22 | 40231 |
| sub-CC00930XX15 | 43930 |
| sub-CC00938XX23 | 47931 |
| sub-CC00939XX24 | 36230 |
| sub-CC00940XX17 | 44030 |
| sub-CC00947XX24 | 43430 |
| sub-CC00948XX25 | 30030 |
| sub-CC00949XX26 | 55630 |
| sub-CC00956XX16 | 37730 |
| sub-CC00958XX18 | 40630 |
| sub-CC00962XX14 | 57830 |
| sub-CC00966XX18 | 46430 |
| sub-CC00967XX19 | 58330 |
| sub-CC00974XX18 | 46931 |
| sub-CC00976XX20 | 64331 |
| sub-CC00980XX16 | 66630 |
| sub-CC00982XX18 | 64930 |
| sub-CC00987XX23 | 66031 |
| sub-CC00990XX18 | 70230 |
| sub-CC00992XX20 | 62530 |
| sub-CC01004XX06 | 51630 |
| sub-CC01007XX09 | 67630 |
| sub-CC01014XX08 | 59530 |
| sub-CC01015XX09 | 74130 |

| | |
|---|---|
| sub-CC01022XX08 | 79430 |
| sub-CC01023XX09 | 74030 |
| sub-CC01027XX13 | 82630 |
| sub-CC01029XX15 | 84830 |
| sub-CC01032XX10 | 77330 |
| sub-CC01037XX15 | 77430 |
| sub-CC01041XX11 | 81131 |
| sub-CC01042XX12 | 74431 |
| sub-CC01044XX14 | 72531 |
| sub-CC01047XX17 | 81830 |
| sub-CC01050XX03 | 82330 |
| sub-CC01051XX04 | 83930 |
| sub-CC01055XX08 | 77830 |
| sub-CC01070XX07 | 86430 |
| sub-CC01074XX11 | 63930 |
| sub-CC01082XX11 | 68830 |
| sub-CC01084XX13 | 69330 |
| sub-CC01086XX15 | 100430 |
| sub-CC01087XX16 | 99430 |
| sub-CC01093AN14 | 75130 |
| sub-CC01093BN14 | 75230 |
| sub-CC01096XX17 | 76130 |
| sub-CC01103XX06 | 101930 |
| sub-CC01117XX12 | 84730 |
| sub-CC01145XX16 | 98330 |
| sub-CC01176XX14 | 132530 |
| sub-CC01190XX12 | 143030 |
| sub-CC01191XX13 | 130330 |
| sub-CC01194XX16 | 149530 |
| sub-CC01195XX17 | 152230 |
| sub-CC01198XX20 | 140930 |
| sub-CC01199XX21 | 141130 |
| sub-CC01200XX04 | 154330 |
| sub-CC01207XX11 | 150330 |
| sub-CC01211XX07 | 145930 |
| sub-CC01215XX11 | 146331 |
| sub-CC01220XX08 | 148731 |
| sub-CC01223XX11 | 149330 |
| sub-CC01236XX16 | 155830 |

Table S3. Longitudinal dataset

| ID | sesion |
|---|---|
| sub-CC00301XX04 | 96400 |
| sub-CC00301XX04 | 113001 |
| sub-CC00305XX08 | 98101 |
| sub-CC00305XX08 | 115700 |
| sub-CC00326XX13 | 104100 |
| sub-CC00326XX13 | 118800 |
| sub-CC00385XX15 | 118500 |
| sub-CC00385XX15 | 125700 |
| sub-CC00389XX19 | 119100 |
| sub-CC00389XX19 | 133800 |

| | |
|---|---|
| sub-CC00529AN18 | 151300 |
| sub-CC00529AN18 | 170000 |
| sub-CC00569XX17 | 158300 |
| sub-CC00569XX17 | 170600 |
| sub-CC00617XX15 | 176500 |
| sub-CC00617XX15 | 188400 |
| sub-CC00632XX14 | 183300 |
| sub-CC00632XX14 | 196000 |
| sub-CC00648XX22 | 191100 |
| sub-CC00648XX22 | 204400 |
| sub-CC00672AN13 | 197601 |
| sub-CC00672AN13 | 214900 |
| sub-CC00712XX11 | 221400 |
| sub-CC00712XX11 | 232701 |
| sub-CC00770XX12 | 238000 |
| sub-CC00770XX12 | 1100 |
| sub-CC00792XX18 | 244200 |
| sub-CC00792XX18 | 1800 |
| sub-CC00823XX15 | 15810 |
| sub-CC00823XX15 | 27810 |
| sub-CC00845AN21 | 26510 |
| sub-CC00845AN21 | 32010 |
| sub-CC00845BN21 | 26410 |
| sub-CC00845BN21 | 32110 |
| sub-CC00855XX14 | 30210 |
| sub-CC00855XX14 | 530 |
| sub-CC00867XX18 | 37111 |
| sub-CC00867XX18 | 8930 |
| sub-CC00889AN24 | 2220 |
| sub-CC00889AN24 | 9230 |
| sub-CC00889BN24 | 2320 |
| sub-CC00889BN24 | 9330 |
| sub-CC00907XX16 | 4230 |
| sub-CC00907XX16 | 19230 |
| sub-CC01005XX07 | 36930 |
| sub-CC01005XX07 | 49930 |
| sub-CC01077XX14 | 65230 |
| sub-CC01077XX14 | 78430 |
| sub-CC01218XX14 | 147430 |
| sub-CC01218XX14 | 157231 |

Table S4. The brain regions showed significant difference between groups for clustering coefficient after FDR correction at q < 0.05.

| ***Regions*** | ***Abbr.*** | ***P-values*** | ***F-values*** | ***Q-values*** |
|---|---|---|---|---|
| Postcentral gyrus right | PoCG-R | 0.000 | 8.349 | 0.000 |
| Amygdala right | AMYG-R | 0.000 | 7.835 | 0.001 |
| Supramarginal gyrus right | SMG-R | 0.000 | 7.766 | 0.001 |
| Precuneus left | PCUN-L | 0.000 | 7.709 | 0.002 |
| Supramarginal gyrus left | SMG-L | 0.000 | 7.555 | 0.002 |
| Paracentral lobule left | PCL-L | 0.000 | 6.855 | 0.003 |
| Superior temporal gyrus right | STG-R | 0.000 | 6.638 | 0.003 |
| Postcentral gyrus left | PoCG-L | 0.000 | 6.484 | 0.004 |

| | | | | |
|---|---|---|---|---|
| Superior parietal gyrus left | SPG-L | 0.000 | 6.333 | 0.005 |
| Inferior frontal gyrus (opercular) right | IFGoperc-R | 0.000 | 6.178 | 0.005 |
| Precentral gyrus left | PreCG-L | 0.000 | 6.132 | 0.006 |
| Inferior parietal lobule left | IPL-L | 0.000 | 5.716 | 0.006 |
| Precuneus right | PCUN-R | 0.000 | 5.554 | 0.007 |
| Superior parietal gyrus right | SPG-R | 0.000 | 5.509 | 0.007 |
| Rolandic operculum left | ROL-L | 0.000 | 5.124 | 0.008 |
| Superior temporal gyrus left | STG-L | 0.000 | 5.026 | 0.008 |
| Orbitofrontal cortex (middle) right | ORBmid-R | 0.000 | 4.851 | 0.009 |
| Paracentral lobule right | PCL-R | 0.000 | 4.788 | 0.01 |
| Superior frontal gyrus (dorsal) right | SFGdor-R | 0.000 | 4.632 | 0.010 |
| Rolandic operculum right | ROL-R | 0.000 | 4.292 | 0.011 |
| Middle cingulate gyrus left | MCG-L | 0.000 | 4.247 | 0.011 |
| Cuneus left | CUN-L | 0.000 | 3.944 | 0.012 |
| Superior occipital gyrus right | SOG-R | 0.000 | 3.933 | 0.012 |
| Anterior cingulate gyrus right | ACG-R | 0.000 | 3.908 | 0.013 |
| Middle cingulate gyrus right | MCG-R | 0.000 | 3.897 | 0.013 |
| Superior occipital gyrus left | SOG-L | 0.000 | 3.835 | 0.014 |
| Anterior cingulate gyrus left | ACG-L | 0.001 | 3.776 | 0.015 |
| Middle occipital gyrus left | MOG-L | 0.001 | 3.729 | 0.015 |
| Angular gyrus right | ANG-R | 0.001 | 3.720 | 0.016 |
| Supplementary motor area left | SMA-L | 0.001 | 3.695 | 0.016 |
| Middle occipital gyrus right | MOG-R | 0.001 | 3.691 | 0.017 |
| Orbitofrontal cortex (medial) right | ORBmed-R | 0.001 | 3.545 | 0.017 |
| Inferior parietal lobule right | IPL-R | 0.003 | 3.279 | 0.018 |
| Inferior frontal gyrus (triangular) right | IFGtriang-R | 0.003 | 3.261 | 0.018 |
| Precentral gyrus right | PreCG-R | 0.004 | 3.210 | 0.019 |
| Heschl gyrus left | HES-L | 0.005 | 3.082 | 0.02 |
| Middle frontal gyrus right | MFG-R | 0.006 | 3.054 | 0.020 |
| Insula right | INS-R | 0.007 | 3.003 | 0.021 |
| Insula left | INS-L | 0.007 | 2.948 | 0.021 |
| Inferior temporal gyrus right | ITG-R | 0.009 | 2.875 | 0.022 |
| Orbitofrontal cortex (medial) left | ORBmed-L | 0.011 | 2.787 | 0.022 |
| Supplementary motor area right | SMA-R | 0.011 | 2.783 | 0.023 |
| Superior frontal gyrus (medial) right | SFGmed-R | 0.011 | 2.772 | 0.023 |
| Middle frontal gyrus left | MFG-L | 0.012 | 2.760 | 0.024 |
| Superior frontal gyrus (medial) left | SFGmed-L | 0.013 | 2.702 | 0.025 |
| Inferior temporal gyrus left | ITG-R | 0.018 | 2.582 | 0.025 |
| Middle temporal gyrus right | MTG-R | 0.019 | 2.542 | 0.026 |
| Lingual gyrus left | LING-L | 0.023 | 2.477 | 0.026 |

Table S5. The brain regions showed significant difference between groups for node strength after FDR correction at q < 0.05.

| ***Regions*** | ***Abbr.*** | ***P-values*** | ***F-values*** | ***Q-values*** |
|---|---|---|---|---|
| Precentral gyrus right | PreCG-R | 0.000 | 10.171 | 0.000 |
| Superior frontal gyrus (dorsal) right | SFGdor-R | 0.000 | 8.614 | 0.001 |
| Precentral gyrus left | PreCG-L | 0.000 | 6.781 | 0.001 |
| Superior frontal gyrus (dorsal) left | SFGdor-L | 0.000 | 6.632 | 0.002 |
| Superior frontal gyrus (medial) right | SFGmed-R | 0.000 | 5.771 | 0.002 |
| ParaHippocampal gyrus left | PHG-L | 0.000 | 5.626 | 0.003 |
| Inferior frontal gyrus (opercular) right | IFGoperc-R | 0.000 | 5.300 | 0.003 |
| Rolandic operculum right | ROL-R | 0.000 | 5.228 | 0.004 |

| | | | | |
|---|---|---|---|---|
| Middle cingulate gyrus left | MCG-L | 0.000 | 5.219 | 0.005 |
| Middle cingulate gyrus right | MCG-R | 0.000 | 5.018 | 0.005 |
| Rolandic operculum left | ROL-L | 0.000 | 4.363 | 0.006 |
| Lingual gyrus right | LING-L | 0.000 | 4.024 | 0.006 |
| Inferior frontal gyrus (opercular) left | IFGoperc-L | 0.000 | 4.010 | 0.007 |
| Supplementary motor area right | SMA-R | 0.000 | 3.993 | 0.007 |
| ParaHippocampal gyrus right | PHG-R | 0.001 | 3.807 | 0.008 |
| Inferior occipital gyrus right | IOG-R | 0.001 | 3.777 | 0.008 |
| Amygdala right | AMYG-R | 0.001 | 3.726 | 0.009 |
| Orbitofrontal cortex (medial) right | ORBmed-R | 0.001 | 3.716 | 0.01 |
| Inferior parietal lobule right | IPL-R | 0.001 | 3.693 | 0.010 |
| Inferior frontal gyrus (triangular) left | IFGtriang-L | 0.001 | 3.680 | 0.011 |
| Middle frontal gyrus right | MFG-R | 0.001 | 3.605 | 0.011 |
| Precuneus right | PCUN-R | 0.002 | 3.538 | 0.012 |
| Supplementary motor area left | SMA-L | 0.002 | 3.518 | 0.012 |
| Inferior frontal gyrus (triangular) right | IFGtriang-R | 0.002 | 3.493 | 0.013 |
| Postcentral gyrus left | PoCG-L | 0.002 | 3.480 | 0.013 |
| Inferior parietal lobule left | IPL-L | 0.002 | 3.420 | 0.014 |
| Precuneus left | PCUN-L | 0.002 | 3.410 | 0.015 |
| Superior frontal gyrus (medial) left | SFGmed-L | 0.003 | 3.367 | 0.015 |
| Anterior cingulate gyrus right | ACG-R | 0.004 | 3.245 | 0.016 |
| Insula left | INS-L | 0.004 | 3.184 | 0.016 |
| Superior parietal gyrus left | SPG-L | 0.005 | 3.131 | 0.017 |
| Postcentral gyrus right | PoCG-R | 0.005 | 3.130 | 0.017 |
| Thalamus right | THA-R | 0.005 | 3.089 | 0.018 |
| Supramarginal gyrus right | SMG-R | 0.007 | 2.976 | 0.018 |
| Hippocampus right | HIP-R | 0.008 | 2.899 | 0.019 |
| Orbitofrontal cortex (medial) left | ORBmed-L | 0.012 | 2.748 | 0.02 |
| Superior parietal gyrus right | SPG-R | 0.018 | 2.579 | 0.020 |
| Lingual gyrus left | LING-L | 0.018 | 2.568 | 0.021 |

Table S6. The brain regions showed significant difference between groups for local efficiency after FDR correction at $q < 0.05$.

| ***Regions*** | ***Abbr.*** | ***P-values*** | ***F-values*** | ***Q-values*** |
|---|---|---|---|---|
| Precuneus left | PCUN-L | 0.000 | 12.206 | 0.000 |
| Postcentral gyrus right | PoCG-R | 0.000 | 11.332 | 0.001 |
| Precentral gyrus left | PreCG-L | 0.000 | 10.794 | 0.001 |
| Supramarginal gyrus right | SMG-R | 0.000 | 10.211 | 0.002 |
| Precuneus right | PCUN-R | 0.000 | 9.836 | 0.002 |
| Postcentral gyrus left | PoCG-L | 0.000 | 9.442 | 0.003 |
| Supramarginal gyrus left | SMG-L | 0.000 | 9.045 | 0.003 |
| Inferior parietal lobule left | IPL-L | 0.000 | 8.944 | 0.004 |
| Superior parietal gyrus left | SPG-L | 0.000 | 8.470 | 0.005 |
| Superior frontal gyrus (dorsal) right | SFGdor-R | 0.000 | 8.403 | 0.005 |
| Superior parietal gyrus right | SPG-R | 0.000 | 8.302 | 0.006 |
| Paracentral lobule left | PCL-L | 0.000 | 8.156 | 0.006 |
| Precentral gyrus right | PreCG-R | 0.000 | 7.917 | 0.007 |
| Amygdala right | AMYG-R | 0.000 | 7.837 | 0.007 |
| Middle cingulate gyrus left | MCG-L | 0.000 | 6.803 | 0.008 |
| Rolandic operculum right | ROL-R | 0.000 | 6.754 | 0.008 |
| Supplementary motor area left | SMA-L | 0.000 | 6.695 | 0.009 |
| Superior temporal gyrus right | STG-R | 0.000 | 6.683 | 0.01 |
| Inferior frontal gyrus (opercular) right | IFGoperc-R | 0.000 | 6.628 | 0.010 |

| | | | | |
|---|---|---|---|---|
| Middle frontal gyrus right | MFG-R | 0.000 | 6.532 | 0.011 |
| Superior occipital gyrus left | SOG-L | 0.000 | 6.417 | 0.011 |
| Superior occipital gyrus right | SOG-R | 0.000 | 6.265 | 0.012 |
| Supplementary motor area right | SMA-R | 0.000 | 6.113 | 0.012 |
| Rolandic operculum left | ROL-L | 0.000 | 6.006 | 0.013 |
| Middle cingulate gyrus right | MCG-R | 0.000 | 5.815 | 0.013 |
| Inferior parietal lobule right | IPL-R | 0.000 | 5.646 | 0.014 |
| Paracentral lobule right | PCL-R | 0.000 | 5.228 | 0.015 |
| Middle frontal gyrus left | MFG-L | 0.000 | 5.155 | 0.015 |
| Angular gyrus right | ANG-R | 0.000 | 5.088 | 0.016 |
| Cuneus left | CUN-L | 0.000 | 5.067 | 0.016 |
| Middle occipital gyrus right | MOG-R | 0.000 | 5.065 | 0.017 |
| Superior frontal gyrus (dorsal) left | SFGdor-L | 0.000 | 4.910 | 0.017 |
| Superior temporal gyrus left | STG-L | 0.000 | 4.834 | 0.018 |
| Orbitofrontal cortex (middle) right | ORBmid-R | 0.000 | 4.793 | 0.018 |
| Middle occipital gyrus left | MOG-L | 0.000 | 4.625 | 0.019 |
| Anterior cingulate gyrus right | ACG-R | 0.000 | 4.165 | 0.02 |
| Inferior frontal gyrus (triangular) right | FGtriang-R | 0.000 | 4.005 | 0.020 |
| Anterior cingulate gyrus left | ACG-L | 0.000 | 3.973 | 0.021 |
| Orbitofrontal cortex (medial) right | ORBmed-R | 0.001 | 3.832 | 0.021 |
| Superior frontal gyrus (medial) right | SFGmed-R | 0.001 | 3.819 | 0.022 |
| Superior frontal gyrus (medial) left | SFGmed-L | 0.001 | 3.659 | 0.022 |
| Insula left | INS-L | 0.002 | 3.400 | 0.023 |
| Inferior temporal gyrus right | ITG-R | 0.003 | 3.246 | 0.023 |
| Heschl gyrus left | HES-L | 0.005 | 3.150 | 0.024 |
| Insula right | INS-R | 0.006 | 3.034 | 0.025 |
| Lingual gyrus left | LING-L | 0.007 | 2.965 | 0.025 |
| Calcarine cortex left | CAL-L | 0.007 | 2.961 | 0.026 |
| Inferior temporal gyrus left | ITG-L | 0.007 | 2.950 | 0.026 |
| Orbitofrontal cortex (medial) left | ORBmed-L | 0.007 | 2.949 | 0.027 |
| Inferior frontal gyrus (triangular) left | FGtriang-L | 0.008 | 2.896 | 0.027 |
| Middle temporal gyrus right | MTG-R | 0.010 | 2.807 | 0.028 |
| Inferior frontal gyrus (opercular) left | IFGoperc-L | 0.013 | 2.722 | 0.028 |
| Cuneus right | CUN-R | 0.014 | 2.688 | 0.029 |
| Middle temporal gyrus left | MTG-L | 0.020 | 2.534 | 0.03 |
